# Multi-modal single-cell profiling reveals transcriptional states and commitment dynamics during sexual reproduction in the diatom *Pseudo-nitzschia multistriata*

**DOI:** 10.64898/2026.03.25.714164

**Authors:** Antonella Ruggiero, Tobias Gerber, Leslie Pan, Francesco Manfellotto, Michael Bonadonna, Diana Ordoñez-Rueda, Isabelle Becher, Frank Stein, Monia T. Russo, Detlev Arendt, Maria Immacolata Ferrante

## Abstract

Diatoms, single-celled photosynthetic eukaryotes, traditionally considered primarily influenced by resource availability, are now recognized to exhibit complex developmental processes reminiscent of those in multicellular organisms. Sexual reproduction is an obligate choice in many species, albeit with variable frequency and efficiency. In order to explore decision making mechanisms and cellular heterogeneity inherent to this specific phase of the diatom life cycle, we applied a multi-modal strategy to follow sexual reproduction events in the model diatom *Pseudo-nitzschia multistriata*: i) we employed image-enabled cell sorting to link sexual stages morphology with gene expression ii) we developed a single-nucleus RNA sequencing assay, and iii) conducted proteomic profiling. Combining phenotypes with single-cell profiling allowed us to identify sexual state-specific transcription programs and different states within morphologically identical parental cells, revealing a widespread commitment of the majority of the population to sexual reproduction. This implies that the bottleneck to successful sexual reproduction in this species can be placed downstream of the signalling phase between mating cells, with additional, yet unidentified, factors influencing mating success at sea.

## Introduction

Oceanic food-webs and the main biogeochemical cycles are based on unicellular eukaryotes belonging to different lineages, characterized by sophisticated morphologies and unique physiological adaptations^1^.Their life cycles, selected over a long evolutionary history, involve distinct phases and stages that have considerable implications for population dynamics. Examples are the bi-phasic life cycle of coccolithophores^2,3^, the formation of dormant resting stages^4,5^ the sexual phase^6,7^, the switch from single cells to colonies^8,9^. While evidence of the ecological role of life cycle transitions is growing in the last decades, the understanding of the molecular mechanisms that regulate them is still at its infancy. Among unicellular eukaryotes diatoms are the most diverse and largest group accounting for approximately 20% of global primary productivity and playing a key role in biogeochemical cycles^10^. They alternate between vegetative growth and an ephemeral/short-lived sexual phase and can enter dormancy through the formation of resting stages as part of their complex life cycles. Transcriptomic approaches carried out in controlled laboratory conditions provided the first insights on the genetic regulation of diatom life cycles with particular focus on sexual reproduction^11^. This is a key phase needed for genetic recombination and for the formation of large-sized cells that counteract the progressive cell miniaturization linked to mitotic divisions^12^. However, different life stages are present along the progression of the sexual phase: cells can undergo distinct fates and the genetic networks underlying fate choices cannot be discriminated with the bulk transcriptomic approaches used so far. Single-cell RNA sequencing (scRNA-seq) enables the reconstruction of complete life cycles in unicellular organisms, charting gene expression changes across successive stages and revealing intermediate or transient cell states shedding light on how individual cells respond to chemical signals and environmental cues^13^. We applied scRNA-seq to capture the transcriptional profiles during sexual reproduction of the marine model diatom *Pseudo-nitzschia multistriata*^14^. This species has a heterothallic mating system and sex occurs when cells under a critical size encounter a partner of the opposite mating type (MT). Only a fraction of the entire population, approximately 20%, completes to meiosis, gametes fuse in two zygotes that form the auxospore, a soft expandable stage deprived of the frustule, within which a large initial cell with a rigid silica cell wall is produced^15^. The onset of the sexual phase is linked to growth arrest regardless of nutrient availability^7^. Bulk RNA-seq studies have shown that parental cells change their transcriptional profile in response to the perception of the mating partner^16^, suggesting a transcriptional signature for cells committed to sex.

We applied single-nuclei RNA sequencing (snRNA-seq) to disentangle the distinct signatures and cell fates during sex in *P multistriata,* coupled to proteomic analyses, adding information on key proteins required during the different phases. This approach allowed to: (i) identify cell state-specific gene expression programs, (ii) detect transitional cell types, (iii) reconstruct developmental trajectories from vegetative cells to auxospore formation and (iv) assign a putative function to genes with poor or no annotation. Our study offers a powerful framework to uncover the ‘decision-making’ processes in marine microeukaryotes. More broadly, establishing a single-cell transcriptomics assay in phytoplankton opens the door to investigate life cycle dynamics and cellular heterogeneity, key drivers of phenotypic and functional diversity in marine ecosystems.

## Results

### Coupling gene expression to cell morphology during sexual reproduction

To capture hidden heterogeneity in the *P. multistriata* population undergoing sexual reproduction, we implemented single cell RNA-seq approaches. To facilitate the cell states assignment, we first established reference gene expression signatures of sexual stages.

Isolating such stages in diatoms is particularly challenging due to their low frequency within populations and the fragility of these cells, as they represent the only cell wall-free, soft-bodied stages. To overcome this limit, we used image-enabled cell sorting^17^ which combines fluorescence-activated cell sorting (FACS) with the detection of cell morphology and subcellular structures. We set up cross cultures of opposite MTs and followed sexual development for 48 hours which correspond to the time point where specific sexual cell types, gametes, zygotes, and early auxospores, are present (Fig. 1a and ^18^). A specific sorting gating strategy enabled to identify and isolate two distinct cell populations: needle-shaped parental cell and round-shaped, sexual cell types (Fig. 1b). The integration of chlorophyll autofluorescence and nuclear staining enhanced specificity by excluding cellular debris and bacteria (Extended Data Fig. 1). Using this approach, we recovered high quality mini-bulk samples consisting of 20-50 cells for each population. When subjecting the bulk transcriptomes to a PCA, we find that the first PC broadly separates the sexual stages from the parental cells supporting the enrichment of different cellular states through the image sorting (Fig. 1c). We next performed a differential gene expression analysis (Wilcoxon Rank Sum test) by comparing the two populations and identified a specific set of genes for each cell population (Fig. 1d). *MRP1*, a *P. multistriata* specific marker which is highly expressed when MT+ cells are exposed to the direct presence or to the growth medium of MT-strains ^16^, ranks among the top markers for parental cells. This enrichment provides independent support for accurate identification of parental cells. To further characterize candidate genes, we examined their predicted protein structures using AlphaFold and performed structural similarity searches with foldseek (Fig. 1d). For the sexual stage specific gene 0000370, the structural comparison identified the fungal ‘Nuclear fusion protein KAR5’ and the plant protein ‘Gamete expressed 1’ (*GEX1*) as closest homolog, leading to the annotation as *GEX1*. *GEX1* is a known marker in *Arabidopsis* gametophyte development and in other organisms ^19,20^ and its presence in the sexual stages population supports effective enrichment for sexual stages. We also inspected *MRP1*, to which we had been unable to assign any putative function due to the lack of clear sequence homologies ^21^. We found that the predicted MRP1 protein consists of a small beta sheet stabilized by 5 alpha helixes (β–α–β–α–β–α–α–β–α), a structural order that matches best with ribosomal S6–like proteins. However, our structural search indicates also a high similarity with the SEA domain, a common extracellular autoproteolytic metazoan domain, which would be consistent with a putative role of this protein in early pheromone signaling.

**Figure 1.**
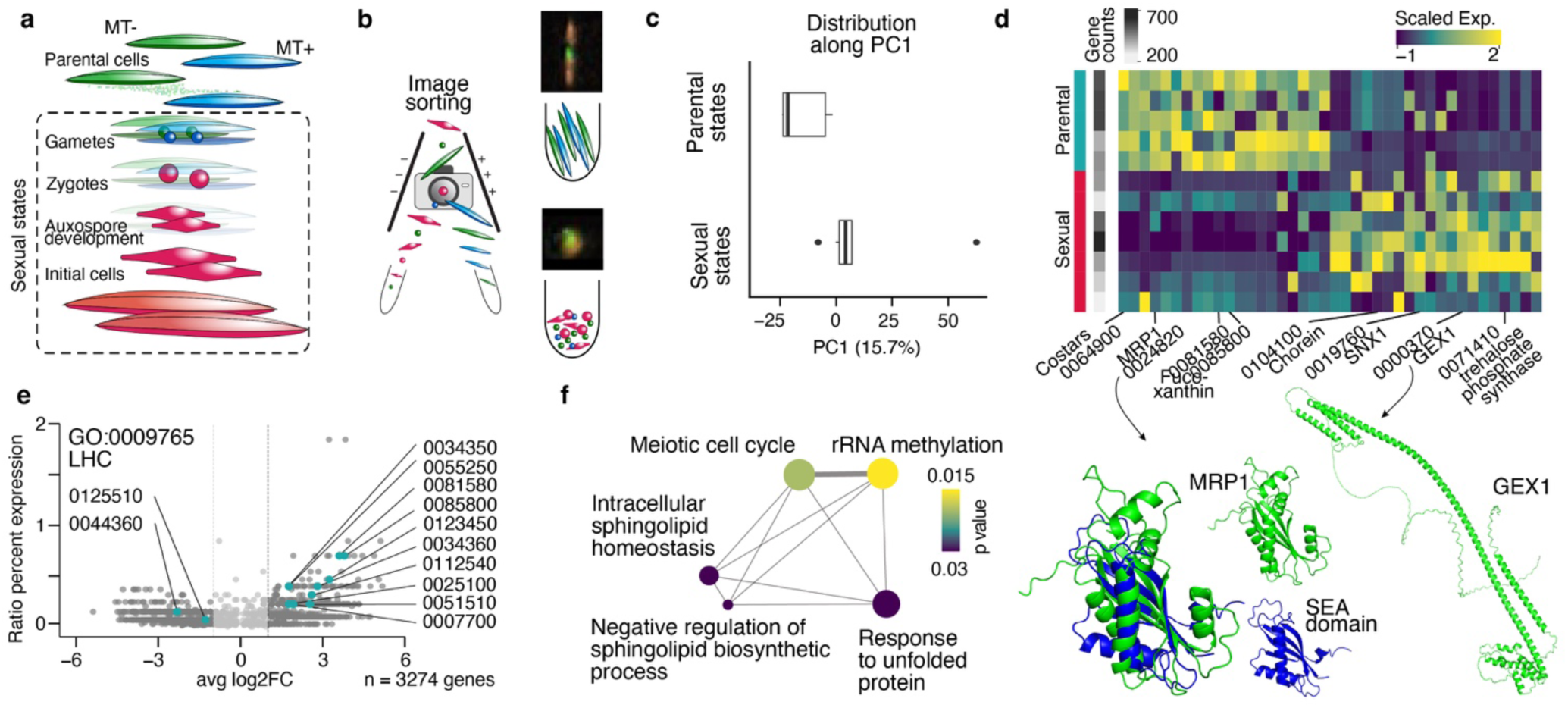
Coupling cell morphology and transcriptomics provides sexual and parental stage specific gene expression. **a** Schematic drawing of the sexual reproduction cycle of *P. multistriata* starting from the pairing of parental cells of opposite MT (blue/green) to the formation of sexual stages and the generation of initial cells (red). **b** Cells were sorted using fluorescence and image derived parameters utilizing an image-based cell sorter. **c** Boxplots showing the distribution of the two populations (parental cell and sexual cell types) across PC1, revealing a separation by transcriptomic signatures. **d** Top: Heatmap showing gene expression of the top 20 markers for each population. Bottom: Predicted protein structures of two state specific markers. Sea domain depicted originates from human MUC1. **e** Volcano plot visualizes all DE genes (grey dots) and highlights genes involved in light harvesting (turquoise, GO:0009765). **f** Network graph shows enriched GO terms for the sexual stages. The color gradient indicates the p-value; circle size indicates the frequency of the GO term in the underlying GOA database. Edge width indicates the degree of similarity.

Top 200 differentially expressed (DE) genes for both populations were subjected to Gene Ontology (GO) enrichment analysis using Goseq. This analysis identified 3 Biological Process (BP) GO terms for the parental population and 5 BP terms for the sexual state population (Fig. 1e-f). For the parental population, the most prominent GO term was Photosynthesis (GO:0009765). Many of the top markers in this population were associated with this function, which was absent from the sexual cell types, suggesting that photosynthesis is reduced during sexual reproduction. However, two genes (0125510 and 0044360) that are also associated with this GO term and that encode for Chlorophyll a-b binding protein L1818 and a Plastid light harvesting protein, respectively, were instead expressed in the sexual fraction (Fig. 1e). GO terms such as Meiotic cell cycle was enriched for the sexual cell types, corroborating the correct identification of genes related to sexual reproduction (Fig. 1f). Among the top markers for the sexual cell types population, we also identified the GO term Response to unfolded proteins (GO:0006986) (Fig. 1f). The expression of stress related genes can be explained by the drastic morphological transitions and the fragility that diatoms face during sexual reproduction, but is also reminiscent of the oxidative burst occurring during fertilization in animals^22^.

Using a morphology-informed sequencing approach, we linked morphological traits to transcriptomic profiles and identified stage-specific gene expression associated with sexual and parental cells.

### Tracking transcriptomic changes during sexual reproduction at single-cell resolution

When cells of opposite MT are mixed, sexual reproduction begins with exchanges of chemical signals and proceeds in a non-synchronous manner over the course of four days. In successful crosses, an obvious growth arrest of the entire cell population until the appearance of the new generation is observed^7,18^. Despite the coordinated response, the number of sexual stages that are actually formed can be very low, and never exceeds 20% of the total cell number. We hypothesized that the parental cells in a cross belong to at least two functionally different populations, one committed to meiosis and the other arrested through unknown mechanisms but not committed. To test this hypothesis, we performed single-nucleus RNA-sequencing (snRNA-seq) experiments along a time course of three days for a single cross using the 10x Genomics platform. We isolated, FACS enriched (Extended Data Fig. 2a) and snap-froze viable nuclei (See Material and Method) to run snRNA-seq experiments on all timepoints in parallel (Fig. 2a). We obtained a total of 1508 nuclei after quality control for data analysis, performed a PCA and reduced the dimensionality by embedding the single-cell transcriptomes by Uniform Manifold Approximation and Projection (UMAP). We identified 7 (c0 - c6) Louvain clusters resembling different cellular states with each of them being defined by a specific set of cluster markers (Table 1, Fig. 2b). We leveraged the morphology based transcriptomic data to score the single-nuclei for the two categories: parental and sexual cell types (Fig. 2b and Extended Data Fig. 2b). The scoring separated out a fraction of cells represented by 2 clusters (c0 and c6) with elevated sexual scores most likely resembling nuclei originating from gametes, zygotes or early auxospores. The majority of nuclei (c1-c5) score higher for the parental state underpinning the generally low fraction of sexual cell types formation.

**Figure 2.**
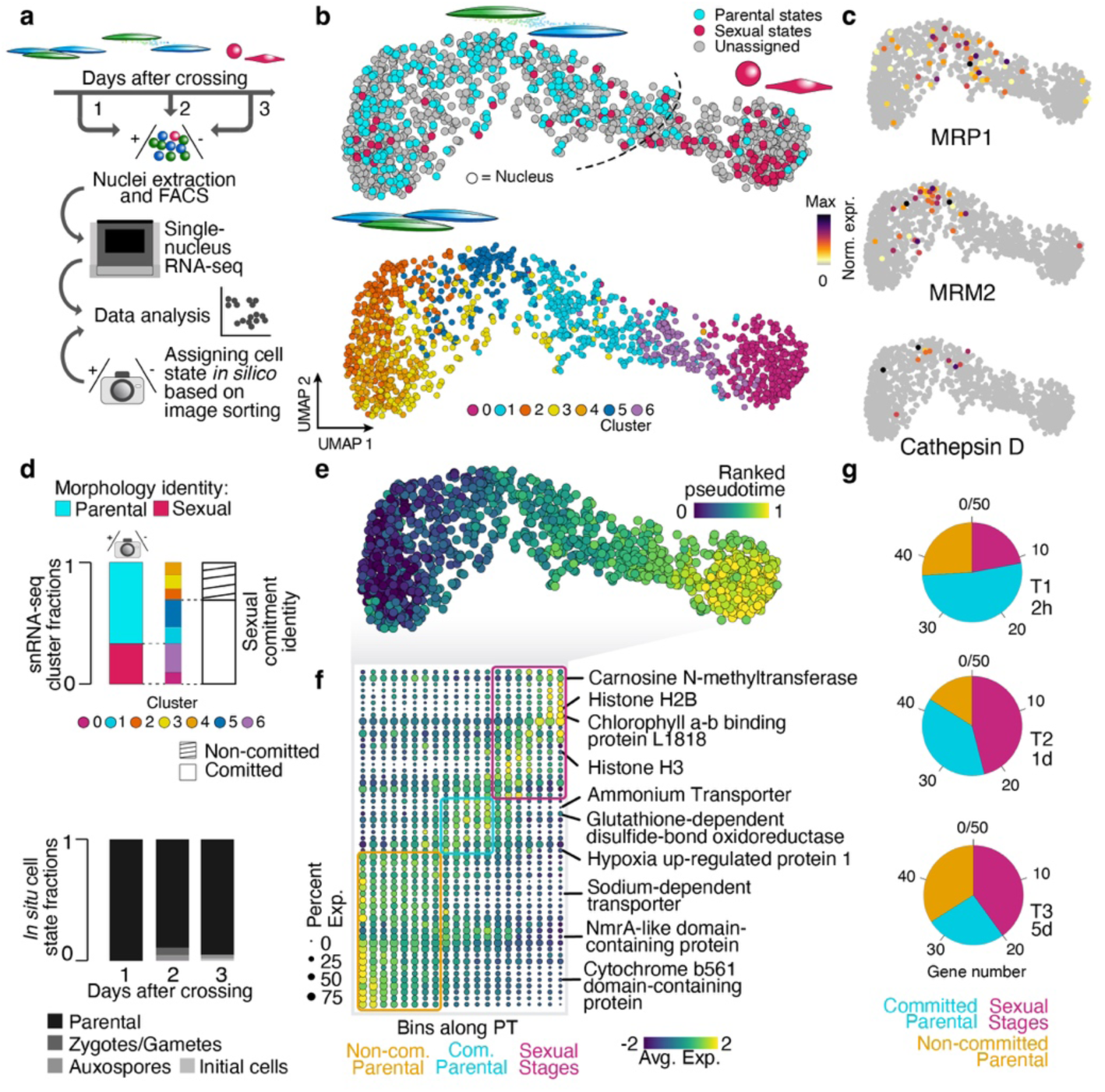
Reconstructing cellular states by single-nuclei transcriptomics reveals a widespread response to mating type crossing. **a** Schematic drawing of the experimental design. SnRNA-seq was performed along a time course of three days. **b** UMAP embeddings visualize cell identities (color-coded) based on morphology scoring for image sorted populations (top) and Louvain clustering (bottom). **c** Plots showing gene expression of three known sexual markers of *P. multistriata* locating MT+ (expressing *MRP1*) and MT-(expressing *MRM2* and cathepsin) committed cells on the embedding. **d** Cellular commitment to the sexual reproduction process. Top: Histogram shows the sexual commitment assessment for scRNA-seq clusters resolved by (left) the morphology scoring alone (parental/sexual) and (right) in combination with marker gene-based cluster annotations. Bottom: Histogram shows the *in vivo* success rate estimated based on the formation frequency of zygotes, auxospores and initial cells assessed with optical microscopy. **e** Pseudotime ordering ranks the nuclei on a developmental trajectory. **f** Gene expression of top 10 cluster markers visualized as dotplot grouped by bins along the pseudotime. Colored boxes indicate group association of marker genes. **g** Top 50 time point specific DE genes of a recent bulk RNA-seq data set across sexual reproduction ^7^(T1:2h, T2:1d, T3:5dpa) are assigned to major single-cell based cell states and fractions are visualized as pie charts. See Extended Data Fig. 2 for the assignment of bulk DE genes to snRNA-seq clusters.

In *P. multistriata* parental cells arrest mitotic division during sexual reproduction with a simultaneous early upregulation of known sexual marker genes as early as 2 hours after crossing cells of opposite MTs^23^. Yet, only a small proportion of cells proceed to gamete formation, leading to a low number of initial cells. To understand the reason for the generally low rate of gamete formation we sought to further annotate the parental clusters c1-c5 in addition to the morphotype scoring. To further resolve the parental clusters, we utilized the expression of known markers such as *MRM2* and cathepsin D, specific for MT-strains, and *MRP1*, specific for MT+ strains, that are upregulated when the other MT is sense^16^. Interestingly, we found one parental fraction corresponding to MT+ (MRP1 positive, c1) and another corresponding to MT-(MRM2 positive, c5) cells that are located adjacent to the sexual stage nuclei (c0/c6), and that we therefore classify as committed to sexual reproduction (Fig. 2c). We consider the remaining and not classified parental populations (c2-c4) as not participating (non-committed) in the sexual reproduction process as they show no sign of neither sexual stage formation nor the expression of key markers for sexual commitment by sensing the opposite MT. Next, we used this classification of clusters to assess the ratio of (i) cells being committed to the sexual reproduction process either by sensing and responding to the opposite MT (c1/c5) or by forming sexual stages (c0/c6) and (ii) cells showing no sign of commitment (c2-c4) (Fig. 2d). Our assignment suggests that on average more than 60% of cells are committed to sexual reproduction in contrast to the frequency of sexual stages observed by microscopy. Next, we calculated a pseudotime trajectory across all nuclei^24^. Pseudotime trajectory analysis reconstructed the transition from the mixed parental populations to sexual cell types by ordering cells along an inferred temporal axis based on transcriptomic similarity. (Fig. 2e). The directionality of the trajectory was confirmed by correlating sexual state scores with increasing pseudo time values and by the lower number of sexual states in the day 1 sample in which the sexual cell types are rare or completely absent (Extended Data Fig. 2c-d). The trajectory allows to visualize genes that gradually change during the course of sexual reproduction (Fig. 2f), corroborating sex specific patterns found in bulk studies such as Chlorophyll a-b binding protein L1818 (0125510)^7^ and the Homologous-pairing protein 2 like domain containing gene (Hop2 like/0010180) ^25^ (Extended Data Fig. 2e). Interestingly we found histone H2B and H3A genes, and genes involved in histone modifications like the Histone Acetyltransferase HAC1^26^, with changing expression patterns towards the sexual cell types population, suggesting chromatin remodeling during this phase of the life cycle (Fig. 2f and marker gene table). Multiple heat shock proteins (Chaperone protein ClpB1, Hypoxia up-regulated protein 1) also change expression patterns across the auxospore formation process (Fig. 2f and marker gene table). Additionally, various transporters like ammonium (0012680) or a sodium dependent transporter (0076160) are differentially expressed across the trajectory. This ammonium transporter is downregulated in bulk data from cells exposed to chemical cues of the opposite MT ^27^ and is part of a family of five members, two of these we also found downregulated during sexual reproduction (0116080 and 0005130). Heterothallic pennate diatom sexual reproduction is not supposed to be dependent on environmental signals, and this downregulation has been interpreted as part of a general reduction of nutrient uptake due to the growth arrest.

To combine cellular resolution with robust gene expression we integrated published *P. multistriata* bulk RNA-seq dataset from a sexual reproduction event ^7,27^ and single-cell transcriptomic data. We compared the transcriptomic patterns of the top 50 time point–specific markers (T1: 2 h, T2: 1 d, T3: 5 d) identified in the bulk time course with their expression profiles in the single-nuclei time course data (Extended Data Fig. 2f). The snRNA-based cell state assignment of the genes highlights a gradual change across the time points with the most T1 (2h) specific genes being expressed by committed parental cells (Fig. 2g). In contrast, T2 (1d) specific genes are more frequently expressed in the sexual cell types. After 5 days (T3) the least genes are assigned to committed cells reflecting a decline in the number of committed cells which aligns with the end of the sexual reproduction phase and appearance of the new generation.

Taken together, this single-cell transcriptomic result enabled us to highlight cellular heterogeneity and to trace a developmental program during the sexual reproduction of *P. multistriata*.

### Genotype assignment resolves non-committed cluster identity

Early decision-making processes that lead to a successful mating occur within the first 24 hours. To capture these events, we performed two additional 10x Genomics snRNA-seq experiments collecting cells at this time point (Fig. 3a and Extended Data Fig. 3a-b). After quality control we retrieved in total 5,373 nuclei for data analysis, integrated the two data sets using Harmony, embedded the data points in a UMAP and clustered the transcriptomes by similarity revealing multiple cellular states (Fig. 3b and Extended Data Fig. 3c-g). We found a previously undetected actively dividing mitotic population identified by known cell cycle markers in diatoms (Extended Data Fig. 3d,e,g) ^28^. This is consistent with the FACS profile, which indicates an incomplete growth arrest with some cells continuing through the cell cycle (Extended Data Fig. 3a-b). We next explored the cell cycle clusters (Fig. 3c and Table 2). In particular, we detected the expression of CyclinD which drives the G1/S progression and CyclinB1-2 which is essential in the G2/M phase (Fig. 3d). The sub-clustering in combination with a pseudotemporal ordering showed a progression through the cell cycle stages (Fig. 3c), providing a *P. multistriata* specific catalogue of cell cycle related genes.

**Figure 3.**
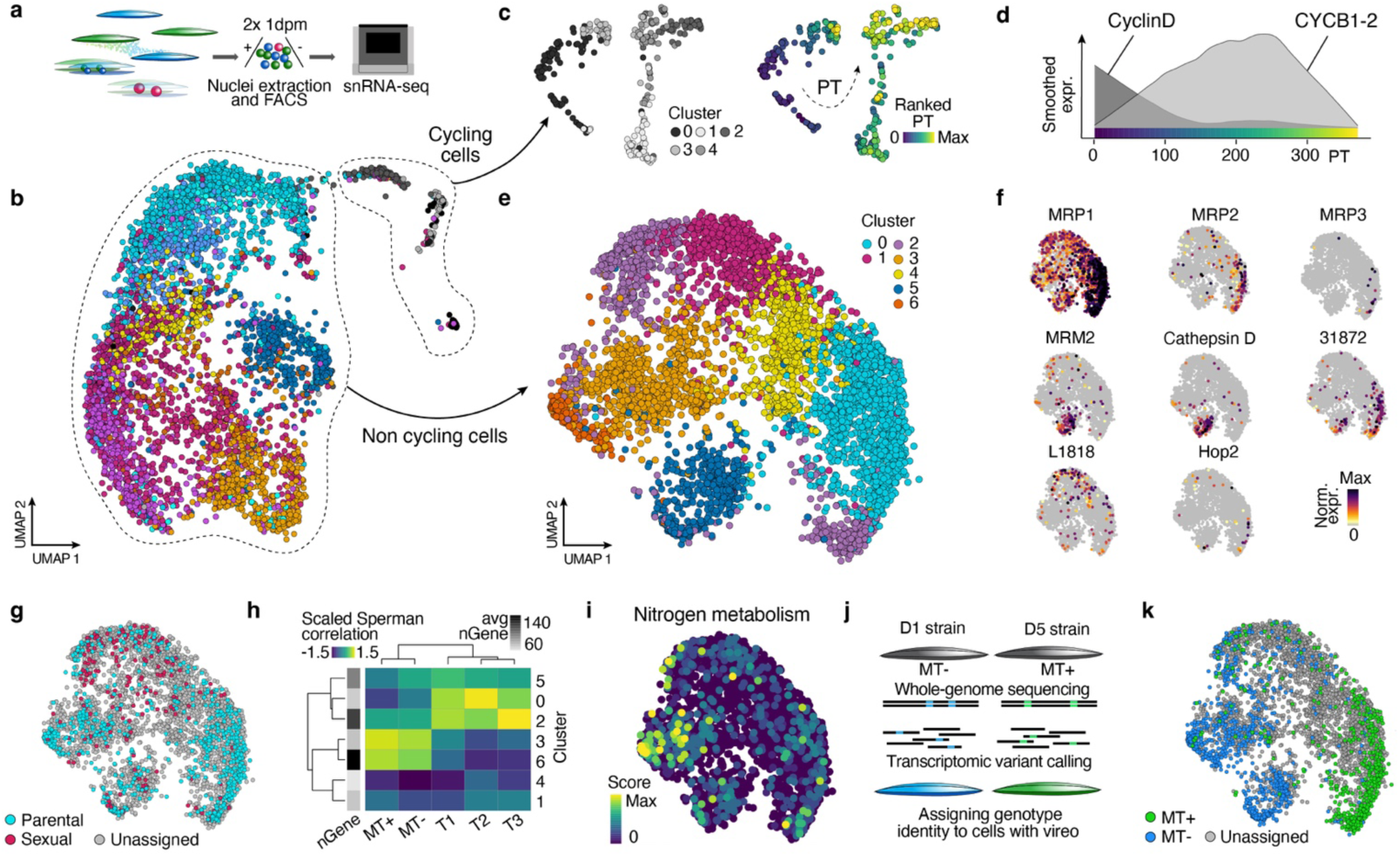
Single-cell genotyping reveals the origin of non-committed cells. **a** Two independent snRNA-seq experiments have been performed on a cross sampled at 24 hrs. **b** Combined UMAP embedding of nuclei of both experiments. Original Louvain cluster colors are applied (see Extended Data Fig. 3). **c** Cycling cells of b are re-embedded and re-clustered (Louvain) (left) and ordered on a pseudotemporal trajectory (right). **d** Ridgeplot showing the early expression of CyclinD and delayed expression of CyclinB1-2 along the pseudo-temporal trajectory. **e** Integrated UMAP embedding showing Louvain cluster identities (color-coded) after removing cycling cells. **f** Feature plots show gene expression of known markers for MT specific committed states and two sexual state specific markers (Chlorophyll a-b binding protein L1818 and Hop 2) identified before (see Extended Data Fig. 1c). **g** Cell identities based on scoring for image sorted populations are shown on the embedding. **h** Heatmap plot shows Spearman correlation between averaged gene expression values per cluster in e with the time course bulk RNA-seq data^7^, T1: 2h, T2: 1d, T3: 5d after crossing. **i** Plot representing a nitrogen metabolism score based on genes identified in^7^. **j** Schematic visualization of the *in silico* genotyping strategy. **k** Genotype identity was assigned based on strain specific SNPs and cellular origins are visualized as features on the embedding.

Next, we removed cycling cells and re-embedded and clustered the data to obtain a data set free of the confounding effect of the cell cycle signal (Fig. 3e). Known marker genes, including previously identified markers (MT+: *MRP1*/*2*/*3*; MT-: *MRM2*/cathepsin D; Sexual states: L1818/Hop2like) (Fig. 3f), together with the scoring for parental and sexual state identities (Fig. 3g and Extended Data Fig. 3h-i) (see also Fig. 1 and 2), allowed us to resolve and disentangle cell states. We found that clusters 1 and 2 represent sexual stages whereas the other clusters represent parental states (Extended Data Fig. 3i). In addition, we validated cell state assignment by correlating averaged cluster transcriptomes to the bulk RNA-seq dataset from^7^. Clusters 0 (committed MT+), 2 (sexual stages) and 5 (committed MT-) showed the highest similarity to the crossed bulk samples whereas non-committed parental clusters of our data set (c3/c6) correlate highest with the uncrossed control samples (Fig. 3h). Clusters 1 and 4 showed no correlation most likely due to a low number of detected genes (Fig. 3h). The non-committed populations showed the highest similarity with the parental monoculture of the bulk RNA-seq indicating that non-committed cells maintain their original (pre-cross) state, while the committed cells undergo a drastic transcriptomic change. Consistent with this, metabolic genes are downregulated in committed parental and sexual cells while remaining active in the non-committed cells (Fig. 3i).

In summary, our classification allowed us to identify a higher number of committed cells (c0/c5) compared to the time course data (Fig 2b), providing an enriched catalogue of shared and MT specific genes for committed parental cells. In order to better characterize the non-committed cells of the two MTs (c3, c4 and c6), we pursued an alternative strategy. We utilized the published genomes of the two strains that were used to perform the experiment, D1 (MT-) and D5 (MT+)^29^, to identify MT unique SNPs^30^ (Fig. 3j). Genetic variation between the parental strains allowed the assignment of the cellular origin to a subset of the single-cell transcriptomes (Fig. 3k and Extended Fig. 3l). The rate of genotype detection is linked to the number of detected genes per nucleus (Extended Data Fig. 3m). Nevertheless, we could validate the correct genotype assignment for the cells with enough genetic information using the MT identity of the committed parental clusters expressing MT specific genes ^31^ (Extended Data Fig. 3l). The *in silico* genotyping revealed that clusters 3 and 6 originate from the D1 MT-strain. For cluster 4 we could not collect sufficient genotype information, however, the transcriptomic similarity suggests that it resembles an MT+ cluster.

Taken together, these analyses enabled the generation of a comprehensive cell state atlas for a crossed culture at 24 hours. We found that the majority of cells sense and respond to the presence of the opposite MT through a drastic reprogramming, whereas a fraction of the population remains unresponsive, maintaining vegetative strain specific characteristics and not engaging in sexual reproduction.

### Tracing zygote formation reveals premeiotic arrest of committed parental cells

We sought to investigate the cellular trajectory for committed cells towards sexual cell states. We isolated the committed and the sexual cell states (cluster 0, 2 and 5) (Extended Data Fig. 4a) and utilized Monocle3 to infer cellular trajectories (Fig. 4a). Starting from the two committed MT specific populations, Monocle3 finds a fusion point within the trajectory, which we interpret as the formation of zygotes. This is supported by the finding, prior to the fusion point, of GO terms related to meiosis, driven by genes like SMC1 (GO:0007064, mitotic sister chromatid cohesion and microtubule process) (Fig. 4b-c and Extended Data Fig. 4b). We therefore explored more canonical diatom meiosis markers^23^ to generate a Meiosis score (Extended Data Fig. 4c). Interestingly, we detected elevated levels of meiosis genes across all committed cells. The score declines towards zygotes and early auxospores (Fig. 4c), which instead are enriched in stress related genes (GO:0009408) (Fig. 4b), in line with other heat stress proteins identified within the time course before (Fig. 2f). The identities of premeiotic committed cells and zygotes/early auxospores are further validated by the reduction in the genotype assignment rate in zygotes besides a high number of detected genes. Genotype assignment is expected to be reduced in zygotes as they comprise a mix of the parental genomic variants and are thus not unambiguously assignable to a single MT (Fig 4d-e). We found no evidence of diatom meiotic genes specific for gametes like the M1 gene^32^, or early prophase markers like Spo11-2 or RAD51a neither within the committed population nor in the zygote cluster. We believe there is a narrow window for the expression of these genes and that our sampling time and the FACS enrichment led to the loss of nuclei from those stages^18^. Nevertheless, these results suggest that committed parental cells are in a pre-meiotic G1 arrest. Consistent with this is that cycling cells originate from non-committed parental cells (Fig. 4f-g and Extended Data Fig. 4d-e) suggesting that committed cells are arrested and do not undergo further mitotic division. Additionally, the non-committed cells show a higher metabolic activity compared to the committed and sexual cells (Fig. 3e). Together, these findings challenge the notion of a dedicated mitotic block underlying the well-characterized growth arrest. Instead, they support a model in which growth arrest arises indirectly as committed cells transition into the meiotic program.

**Figure 4.**
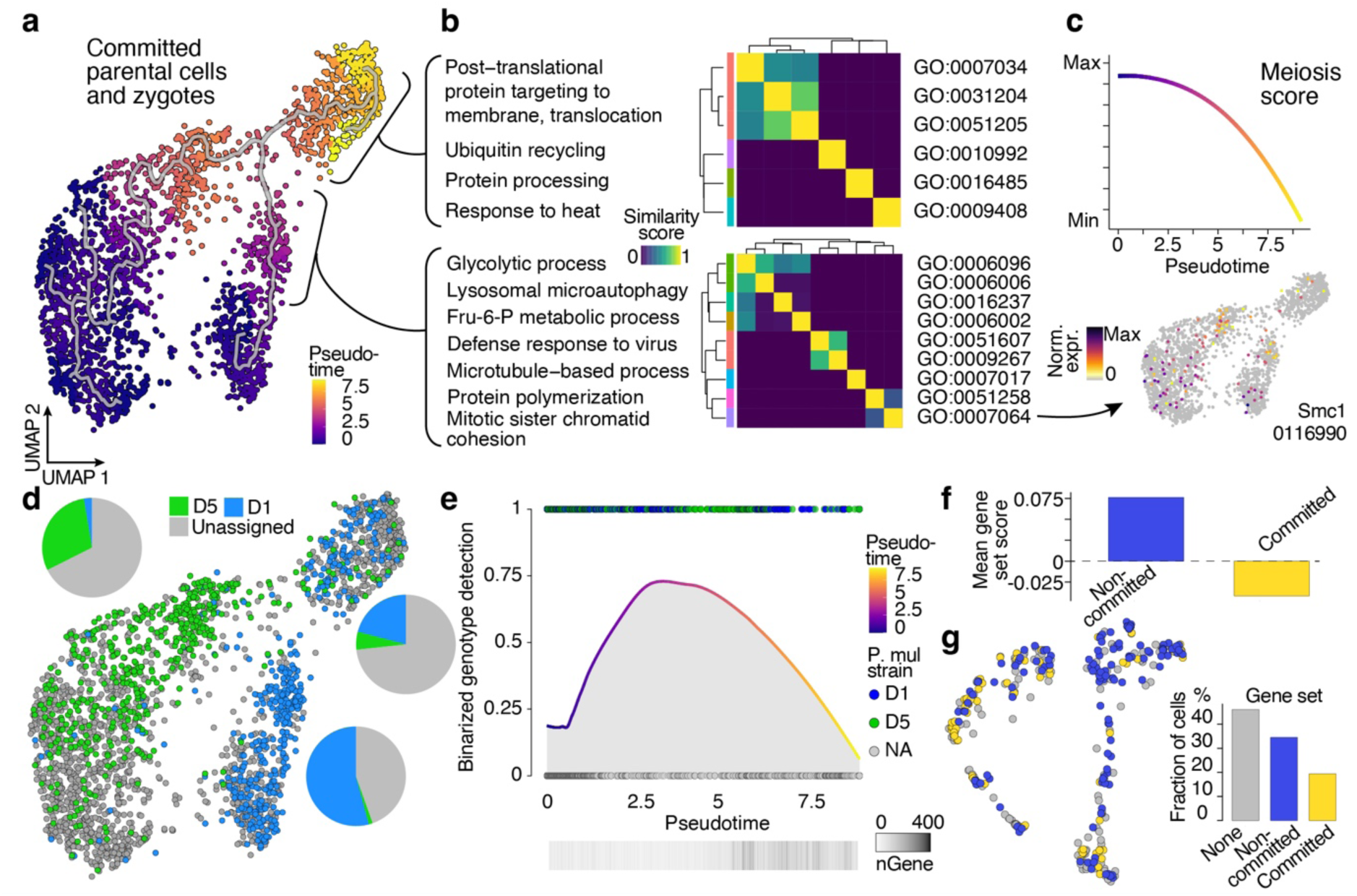
Cell commitment initiates the meiotic program arresting cells in G1. **a** UMAP embedding of committed MT-(cluster 5) and MT+ (cluster 0) and early sexual state cells (cluster 2) (see Fig. 3b). A trajectory was estimated by Monocle3 starting from committed cells and fusing towards zygotes; cells are ordered by pseudotime. **b** The result of a GO enrichment analysis performed on markers before and after the fusion point, respectively, is visualized by the GO term similarities as heatmap. **c** Top: A meiosis score was calculated across all cells and is plotted along pseudotime. Bottom: The expression of Smc1, a cohesin component associated with GO:0007064, is plotted on the embedding. **d** Strain origin predictions are visualized on the embedding and fractions per group (committed MT+/- and zygotes) are shown as pie charts. **e** Smoothed Loess function through the binarized (0 or 1) genotype assignments for cells along pseudotime. Reduced genotype assignment in zygotes is independent of gene detection rates (bottom). **f** Cycling cells were scored for two genesets: non-committed and committed states and the averaged scores are shown as histogram. **g** Scores in f were used to assign identities to the cycling cells. Histogram shows the fraction of each identity across all cells with the non-committed identity being twice as frequent as the committed identity.

In line with the notion that transitions between mitotic and meiotic states in eukaryotes are largely governed by phosphorylation of key regulatory factors^33,34^, we generated a phosphoproteomic dataset (Table 4) from monoclonal parental diatom cells under naive conditions to explore potential post-translational regulation of cell cycle-associated proteins. Notably, phosphorylation was detected on an RNA-binding protein (RBP), the serine/threonine kinase MPS1, and components associated with the mTOR pathway. These proteins are central to processes including checkpoint control, spindle assembly, and nutrient-dependent growth regulation.

### Proteomics data extends the multimodal atlas of cellular states during sexual reproduction in diatoms

We produced a comprehensive proteomic study during distinct sexual reproduction phases in *P. multistriata*. We set up duplicate crosses of D1 (MT-) and D5 (MT+) and sampled them after 2 hours, 1 day and 5 days. In addition, we collected controls of the parental monocultures at T0 and after 5 days (Fig. 5a). We identified 3593 predicted proteins which accounted for ∼29.6% of all predicted protein-coding genes (Extended Data Table 1). PCA analysis reveals that only 3% of variation is defined by differences among replicates. (Extended Data Fig. 5a). The largest number of proteins with significant fold change values was found in parental monoculture control samples after 5 days when compared to T0 (Extended Data Table 1). We performed a GO enrichment analysis on the significantly up and down regulated proteins in this comparison and we found that the biological processes (BP) glucose and carbohydrate metabolic processes have the highest significance among the up regulated proteins whereas the BP of photosynthesis is among the top terms in the downregulated proteins (Fig. 5b-c). We attribute these changes to the switch from exponential to stationary growth. Similarly, when comparing across conditions along the time course, the largest difference was observed at the last time point suggesting that major changes in protein composition are linked to physiology and not specifically to the sexual reproduction ongoing in the crossed culture. Indeed, when comparing the lists of up- and downregulated proteins, there is a strong overlap between the control and crossed cultures, with only a small subset of proteins being sexual reproduction-specific across all comparisons (44 Up/14 Down). To focus on proteins linked to sex, we removed proteins associated with general physiological responses and only considered proteins that changed over the course of the mating event by comparing each time point to the parental cultures (Fig. 5d).

**Figure 5.**
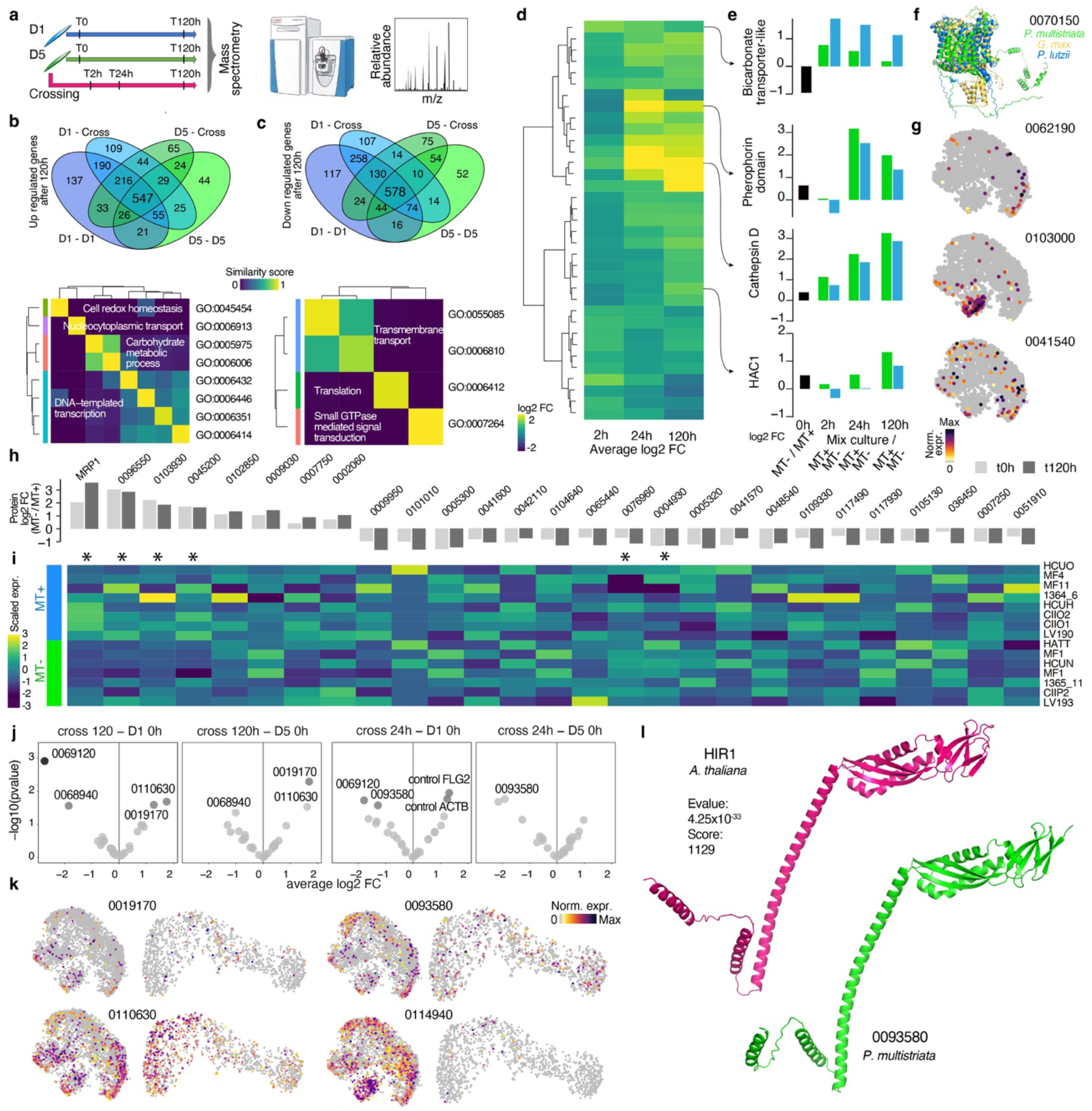
Proteomics analysis identifies key protein changes during sexual reproduction and between mating types. **a** Schematic of the experimental design. **b** Venn diagram (top) shows overlap of significantly up-regulated proteins compared to the initial parental culture after 5 days in monocultures or a cross culture of D1 and D5 strains, respectively. Bottom: GO term enrichment for upregulated genes in the monoculture of D1 or D5. Semantic GO term similarity is visualized as heatmap. **c** The same as in b but for significantly down-regulated proteins. **d** Heatmap shows significantly changing averaged protein expression patterns along the time course of sexual reproduction. **e** Histograms highlight the exact fold changes for certain proteins. **f** Overlay of bicarbonate transporter domain proteins of *P. multistriata*, soy and a fungus. **g** Mapping the gene expression of identified proteins on the snRNA-seq data reveals sex specific expression masked by the bulk proteomics. **h** Fold change values (log2) of consistently differing mating type specific proteins at T0 and T120h in the parental strains are represented as histograms. **i** Proteins in g were screened for gene expression changes across independent strains by a heatmap visualization. Strains represent different MTs. Asterisks mark potential MT specific proteins after evaluating their correlation with a MT specific pseudo gene, respectively. See Extended Data Fig. 5d. **j** Vulcano plots highlight significantly differing secreted proteins between the conditions tested. Significant hits (black), candidates (darkgrey) and selected non-significant proteins (lightgrey) are labelled by name. **k** Feature plots of gene expression of identified proteins in j reveal cell state specific expressions. **l** Aligned predicted protein structures of HIR1 and the *P. multistriata* homolog.

We found a few proteins upregulated as early as 2 hours after crossing, indicative of a fast protein translation changes following mate perception (Fig. 5d-f). One of them is a bicarbonate transporter-like transmembrane domain-containing protein. Nevertheless, protein modelling reveals a high degree of similarity across diatoms, plants and fungi (Fig. 5f and Extended Data Fig. 5c). We also found known genes involved in sexual reproduction like the MT-specific protein CathepsinD^16^ (Fig. 5d-e). In addition, a few other MT specific candidates like a presumably secreted protein (0062190) containing a pherophorin domain is highly upregulated at 1 day after crossing (Fig. 5e) and is specifically expressed in committed MT+ in the single-cell dataset (Fig. 5g). Another group of proteins like HAC1 with a peak in fold change after 5 days of crossing are expressed in auxospores (c1 and c2) of the sn data (Fig. 3e and Fig. 5g). Combining the proteomics with the single-cell data allowed us not only to obtain insights on protein changes but also to assign the cell states in which the proteins are expressed.

Next, we screened for MT specific proteins by leveraging the control samples of the monoclonal parental cultures at 0 h and after 120 h assuming that truly MT specific proteins should not be affected by the physiological differences we have identified before. Our filtering rationale was that a MT specific protein significantly differentially expressed in one of the time points (0 h/120 h) needs to have at least the same trend in the other time point. This screen identified 8 MT+ specific and 19 MT-specific proteins (Fig. 5h). MRP1, one of the best described MT+ specific genes, was among the top hits on the protein level. To refine our list of candidates further, we investigated multiple published bulk RNA-seq studies and assessed the MT specific protein coding gene expression across independent *P. multistriata* strains comprising different MTs. To quantify the similarity, we correlated each gene with a pseudo gene specific to small MT+ cells and found that 4 MT+ specific (including MRP1) and 2 MT-specific proteins remained (r: > 0.35) (Fig. 5i and Extended Data Fig. 5d).

The mate perception signaling and how the planktonic, non-motile *P. multistriata* cells of the opposite MT find each other is still an open question. To explore whether secreted proteins could play a role in these processes we performed a secretomics experiment that followed the experimental design of the proteomics study (Fig. 5a) for which we isolated proteins from the supernatant of the parental cultures and from the cross. One replicate sample fell outside of the variation of the other samples reflecting the difficulties to extract enough protein material from the sea water medium (see methods) (Extended Data Fig. 5a and Extended Data Table 2). After correcting for this batch effect, we detected even fewer proteins (in total 20) of which only 8 also showed a predicted signal peptide sequence (Extended Data Table 2). We attribute the reduced number in conventional signaling peptide sequences in the secreted proteins to the existence of unconventional secretion pathways omitting the ER transport in diatoms^35^ and to possible contamination of highly translated proteins originating from small cell debris. As such, we consider a translation elongation factor (0068940) and a ribosomal protein (0004670) as contamination and ignore them further. Nevertheless, common patterns emerged when comparing the cross culture against the parental cultures prior to the cross. We find that the proteins 0110630 and 019170 are specific to the T0 parental cultures suggesting that the protein expression is downregulated in the crossed culture (Fig. 5j). Indeed, this pattern is consistent with both single-cell data sets that show that zygotes and early auxospores do not express these genes (Fig. 5k). An example for an upregulation during the cross is protein 093580, specific to the 1 day cross when compared to the original cultures (Fig 5j). The single-cell data suggests that this increase in protein abundance is due to a more frequent expression in zygotes and auxospores (Fig. 5k). When we compared the protein structure against databases, we found a very strong agreement with the Hypersensitive-induced response (HIR) protein family of plants (Fig. 5l). In plants it has been shown that HIR1 is associated with lipid rafts which are sterol-and sphingolipid-enriched membrane signaling domains^36^. HIR1 is upregulated in plants after a bacterial treatment suggesting a role in guiding an initial immune reaction through the establishment of lipid rafts. The exact function of HIR proteins during the initial phase of sexual reproduction in *P. multistriata* needs to be determined but our data suggests that the membrane might get actively remodeled to allow a progression during the sexual reproduction cycle. Interestingly, the secretomics data set revealed a D5 strain (MT+) specific secreted protein (069120) (Extended Data Fig. 5b), encoded by a gene which was recently identified among highly methylated *P. multistriata* genes^37^. To identify MT specific proteins that could complement the genes identified in transcriptomic analyses will be essential to expand the proteomic datasets.

### A triplet domain silica transporter is sexual stage specific

A distinguishing feature of diatoms is their silica shell, the frustule. To form the frustule, diatoms absorb dissolved silicic acid (Si(OH)₄) from seawater, which can either passively diffuse into the cell or be actively transported via silicon transporters (SITs)^38^ (Fig. 6a). We detected a silica transporter (10530) among the top genes expressed in cycling cells (Fig. 6b and Extended Data Fig. 6a-b) as shown for other SIT transporters in the centric diatom *T. pseudonana*. Next, we searched the *P. multistriata* genome for SITs by sequence alignment and identified 5 SITs with clear structural homology to SIT1 of *T. pseudonana* (Extended Data Fig. 6a). Of these, 3 SITs were reliably detected in the single-cell data (Fig. 6c and Extended Data Fig. 6b). SIT 32110 was found to be expressed in cycling cells, while SIT 52940 was more specific to the non-cycling fractions (Fig. 6b-c and Extended Data Fig. 6b). Strikingly SIT 52940 is expressed only by the sexual state (zygotes and early auxospores), suggesting a state specific expression for SITs, with this gene being expressed during sexual reproduction and 32110/10530 expressed in the cycling cell population. When looking at the structure, the cell-cycle-specific SITs consist of a monomer (Fig. 6d) while the sex-specific SIT comprises a triplet of SIT domains. When aligning predicted protein structures of SIT monomers (10530 *P. multistriata* / SIT1 *T. pseudonana* / SIT3 *P. tricornutum*) with the triplet SIT (52940) the monomers and individual triplet domain structures show a high degree of similarity. In addition, there seems to be a higher similarity for two of the triplet domains.

**Figure 6.**
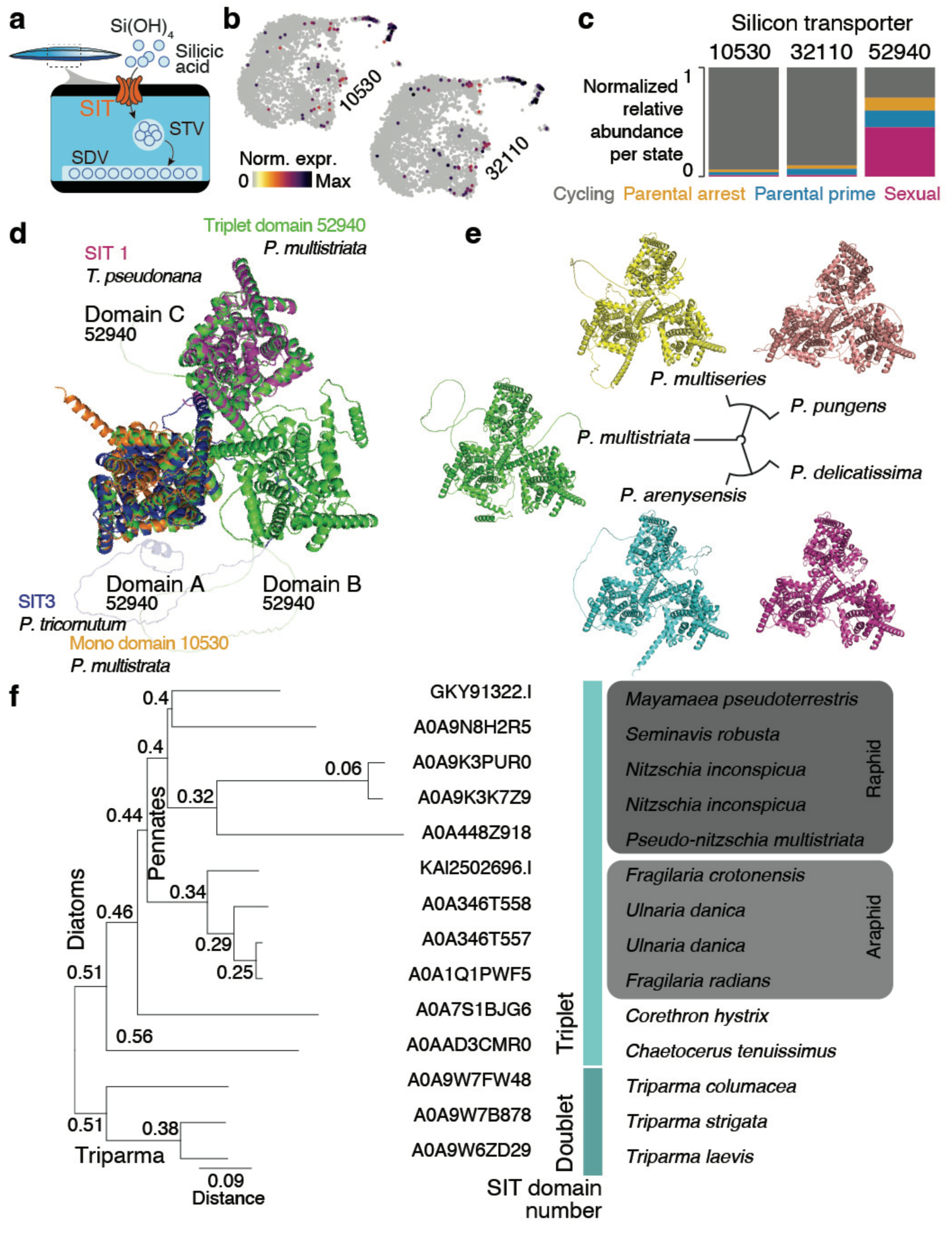
Silica transporters are cell type specifically expressed and their protein structures are evolutionarily conserved. **a** Cartoon of the simplified process of silicic acid uptake through silica transporters (SIT) and the biomineralization and storage process using silica transport (STV) and deposition (SDV) vesicles. **b** Feature plots show gene expression of two SIT mono domain genes specifically in cycling cells after 1 day of crossing (see Fig. 3). **c** Relative binarized gene abundance of the three SITs detected is visualized as histogram for each state after 1 day of crossing. **d** Alphafold prediction for gene 0052940 (green) reveals a triplet domain structure with three SIT transporters in a triangular orientation within the membrane. Mono domain SIT structures of other diatom species with the triplet SIT (color-coded) are aligned. **e** Predicted protein structures for published transcripts of the *Pseudo-nitzschia* genus are aligned and clustered (dendrogram) by their protein sequence alignment. **f** Searching Uniprot revealed triplet domain SITs in 11 diatom species and doublet domain SITs in the diatom sister clade *Triparma*. Triplets were found across all major diatom groups besides *Thallasiosirales*.

Interestingly, a recent study characterizing a monomeric SIT by size-exclusion chromatography suggested that SITs might self-assemble as triplets in the membranes of diatoms as the measured size for *P. tricornutum* SIT1 was three times higher than expected for a SIT monomer ^39^. We therefore modelled the protein interactions to evaluate whether homotrimer formation of our cell cycle specific SIT monomers is possible. Strikingly, Alphafold3 predictions model a homo-trimer protein structure for both cell-cycle specific SITs (52940/10530). In particular, the triplet of the 10530 SIT shows sufficiently high confidence values (ipTM=0.7, pTM=0.73) suggesting a correct prediction of the assembly (Extended Data Fig. 6c-d) with neutral electrostatic charges towards the outside, a positively charged area in between the monomers, 3 negatively charged translocation pathways and a negatively charged ring at the cytosolic surface of the predicted protein structure (Extended Data Fig. 6c-e).

To assess the conservation and hence importance of the triplet domain SIT within the genus of *Pseudo-nitzschia*, we screened published transcriptomes^40^ and predicted protein structures. In all five species: *P. multistriata, P. arenysensis*, *P. delicatissima, P. multiseries* and *P. pungens*, a triplet SIT was identified and the alignment of the sequences and structures highlights a high degree of similarity (Fig. 6e).

Motivated by this, we carried out a survey of triplet SITs across all diatoms in the Uniprot database which revealed the existence of triplet SITs across all major diatom lineages extending a recent study on SIT multimers in diatoms ^41^. In addition, our survey reveals the existence of SIT multimers in the diatom sister clade *Triparma* suggesting that the last common ancestor of diatoms and *Triparma* had already a SIT multimer encoded (Fig. 6f). The phylogenetic tree of multimer SITs resembles in large parts the current phylogeny of diatoms^42^, with minor differences for the position of *Chaetoceros* and *Corethron* species. Interestingly, the only major diatom clade without a SIT triplet besides having many genomics resources available remains the clade of *Thalassiosirales*. Whether the use of a triplet SIT is linked to the occurrence of sexual reproduction in diatoms appears to be an interesting question to follow up on.

## Discussion

In this study, we combined image-based cell sorting with single-nucleus RNA sequencing, proteomics, and structural prediction to resolve, at high resolution, the decision making and developmental trajectories underlying the sexual phase of the marine diatom *Pseudo-nitzschia multistriata*. This integrated approach enabled the molecular definition of discrete sub-populations, including vegetative cycling cells, sexually committed cells, meiotic cells undergoing growth arrest, and sexual stages such as zygotes and auxospores. By directly linking cellular morphology to gene expression and protein profiles, we uncover extensive heterogeneity within isogenic populations and define a continuum of cell states that establishes a temporal framework for sexual development.

These findings advance the mechanistic understanding of sexual reproduction in unicellular eukaryotes, a process resolved in only a few model systems^43,44^. In diatoms, limited genetic and in *situ* approaches have constrained the direct association between morphology and molecular state. Our results pave the way to the elucidation of the molecular and biochemical networks that regulate the life of a unicellular microalga in a way that parallels the development and differentiation of multicellular organisms.

Single-cell analyses revealed substantial transcriptional remodeling during the transition to sexual reproduction, both at the level of parental cells and during progression toward sexual stages. Among morphologically indistinguishable parental cells, we identified multiple transcriptionally distinct states corresponding to the two mating types and to cells committed to sexual reproduction. Remarkably, a large fraction of parental cells appeared committed to sex under mating-compatible conditions, suggesting that sexual readiness may be a prevalent strategy rather than restricted to a rare subset of responsive cells. Following patterns of orientation of pennate diatoms during sinking, it has been proposed that planktonic cells need to sink to stable environments where encounter rates and pairing can occur^45^. From an ecological perspective, the widespread commitment supported by our data is consistent with the low probability of successful mate encounters in the open water column.

We further identified a set of genes associated with the transition toward auxospore formation and defined a meiotic scoring framework that captures progression through gametogenesis and zygote formation. This analysis indicates that committed parental cells broadly express meiotic programs^23^ but experienced a growth arrest until successful mating occurs. These findings support a model in which commitment to sexual reproduction is coupled to a reversible growth arrest, potentially linked to G1 phase regulation (Fig. 7). In contrast, non-committed cells constitute the actively cycling fraction of the population, suggesting that they are not subject to the same arrest program. When mating fails after meiotic entry, we propose that cells may escape checkpoint arrest and revert to mitotic cycling through a mechanism analogous to meiotic slippage described in other eukaryotes ^46^. The molecular basis and regulation of this arrest release system remain to be elucidated.

**Figure 7.**
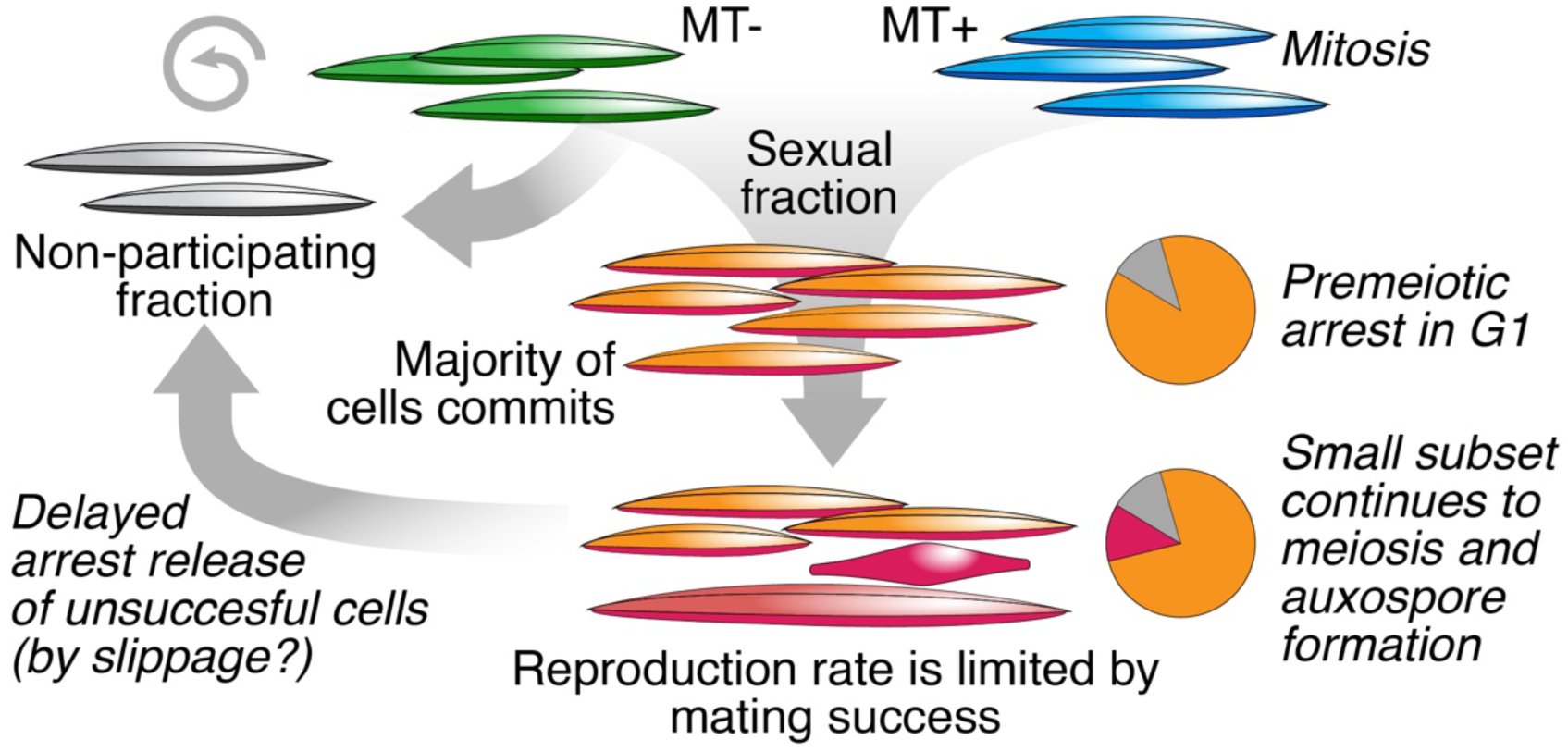
New proposed model of arrest establishment during sexual reproduction.

Our single-cell resource enables a refined, cell-resolved reinterpretation of previously described molecular markers, illustrating the limitations of bulk transcriptomic approaches. A representative case is provided by silicic acid transporters (SITs), which have been reported as downregulated during sexual reproduction based on bulk measurements^7^. In this context, this apparent downregulation has been interpreted as a coordinated transcriptional repression across the population. However, our single-cell analysis reveals that SIT expression is not uniformly reduced across all cells but is instead restricted to a specific subset of cycling cells. Importantly, single-cell resolution further uncovers SIT expression within sexual cell types, suggesting that these transporters may also play roles beyond vegetative growth. The presence of SIT transcripts in defined sexual cell states raises the possibility that distinct structural or regulatory configurations of these transporters are involved in frustule formation during the development of initial cells.

Finally, proteomic and secretomic analyses provide an additional layer of insight into the relationship between gene expression and cellular function during sexual reproduction. While global physiological changes influence the overall protein landscape, the mating process itself appears robust to such variation. Proteomic profiling validated known sexual markers and uncovered additional proteins associated with sexual states, including factors not readily detectable at the transcript level. The inclusion of secreted proteins further offers a direct window into extracellular interactions between mating types. We did not identify canonical sex pheromones in *P. multistriata*, but existing ^16^ and our data provide indirect evidence supporting the existence of chemical communication between mating types, highlighting candidate molecules potentially involved in cell–cell recognition or communication.

Together, these findings establish a multi-modal framework for dissecting cellular heterogeneity and developmental trajectories during diatom sexual reproduction. They reveal that sexual commitment is widespread, transcriptionally primed, and associated with a regulated pre-meiotic arrest, while also pointing to unresolved aspects of mating success, checkpoint control, and environmental modulation.

Recent advances in imaging,^47^ epigenomics^37^, and functional genomics^48,49^ are expected to further extend these insights, enabling a deeper understanding of the molecular logic governing diatom life cycles. Coupling single-cell transcriptomics with the use of transgenic reporter lines for the validation of key genes, as done for another model diatom (Bilcke et al, in preparation), also appears to be a promising avenue for diatom biology. More broadly, this work opens new horizons to explore how unicellular eukaryotes coordinate complex developmental programs that parallel aspects of multicellular differentiation, with implications for cell biology, evolution, and marine ecology.

---

**Extended Data Fig. 1.**
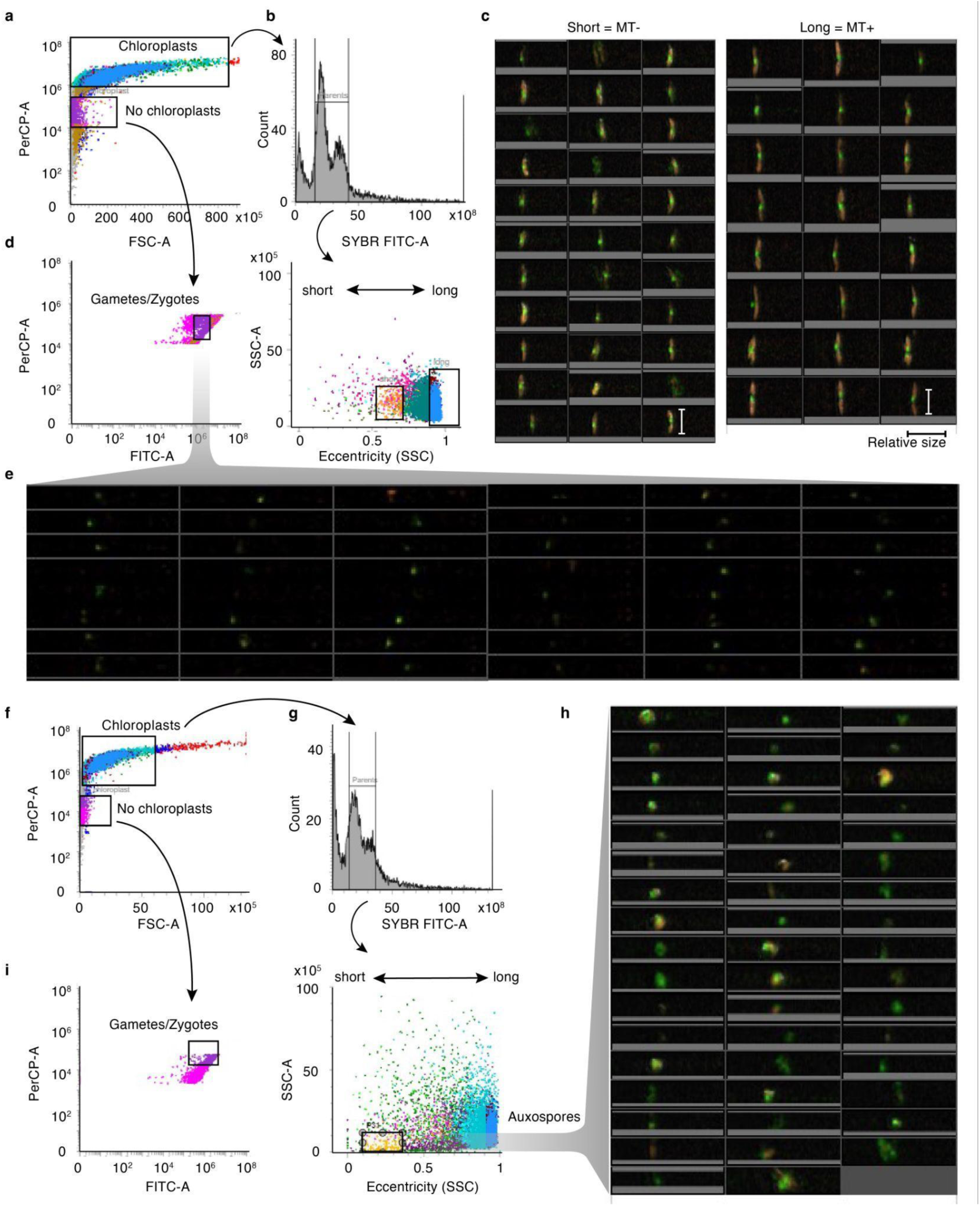
Morphology based cell sorting enriches phenotypes of interest. a-e. FACS image sorting strategy for obtaining different populations of single cells 1 day after crossing strains. **a** The BD image sorter was used to separate populations based on chloroplast autofluorescence. **b** Top: Sybr green signal intensity for chloroplast positive population identifies 2 peaks: one for non-cycling and another for cycling cells exhibiting higher DNA content. Bottom: The chloroplast positive population can be further separated in a longer and a shorter fraction of cells. **c** Virtual exemplary cells are visualized for the long and the short fraction as gated in b representing parental cells of the 2 strains used. **d** Chloroplast negative cells of a are depicted. **e** Exemplary virtual cells are shown for the gate in d. **f-i** The same strategy as in a-e but for the same cross after 2 days. Chloroplast signal is unchanged but the fraction of cycling cells has decreased and among the chloroplast positive fraction a very short auxospore fraction emerged.

**Extended Data Fig. 2.**
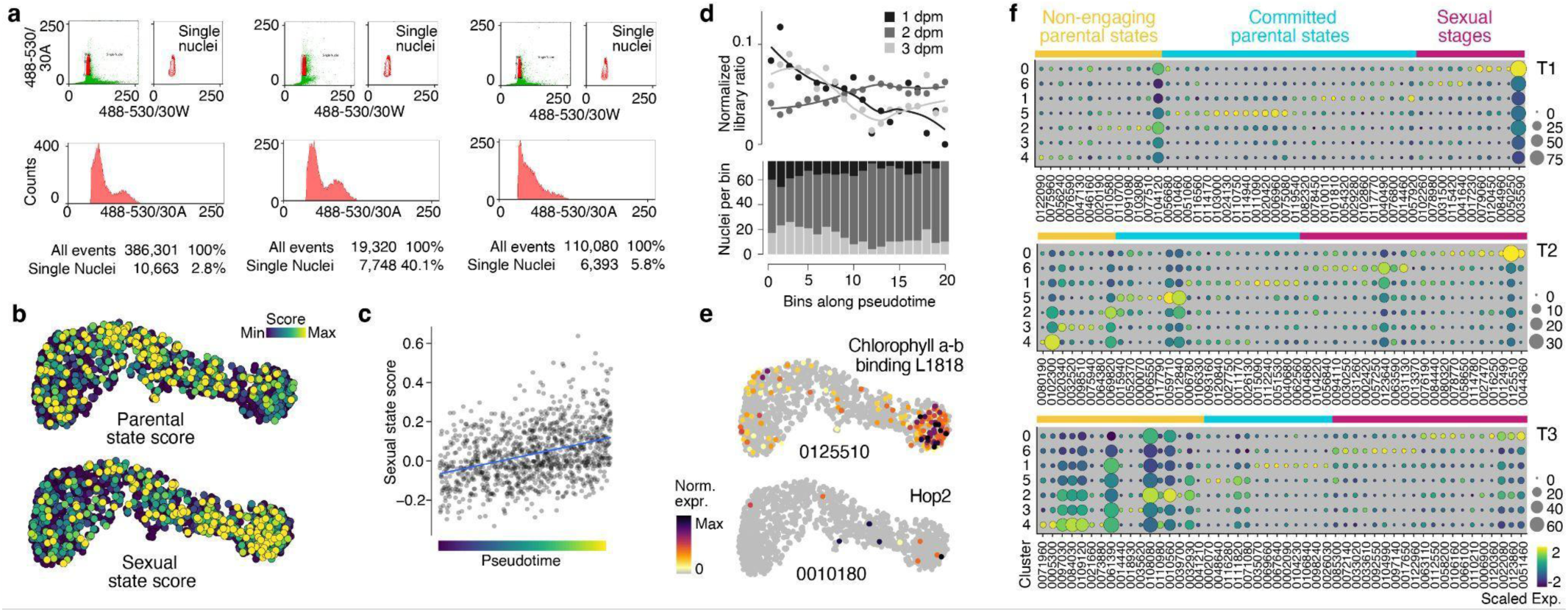
Identifying cellular states during sexual reproduction. **a** FACS profiles applied for separating single nuclei from debris after nuclei extraction for the time course dataset. **b** Scores are calculated based on transcriptomic profiles of the image sorted populations (see Fig. 1) and are visualized on the UMAP. **c** Gene expression of two known sexual state specific markers are shown as features on the UMAP in Fig. 2b **d** Sexual state score positively correlates with pseudo order shown as scatter plot. **e** Contribution of nuclei across different time points along a binned pseudo time trajectory (see Fig. 2b) reveals a decreased number of nuclei for higher pseudo time ranks in the 1 dpm sample. Top: Scatter plot showing group size normalized nuclei fraction per bin. Loess smoothing was applied to fit a line through the data points. Bottom: Histogram with actual nuclei numbers per bin. dpm: days post mating. **f** Top 50 timepoint markers of a bulk RNA-seq study of Annunziata et al.,^7^are visualized by cluster averaged values as dot plot. Assignment to cellular states (colored bars at the top of each dot plot) occurred manually based on cluster expression.

**Extended Data Fig. 3.**
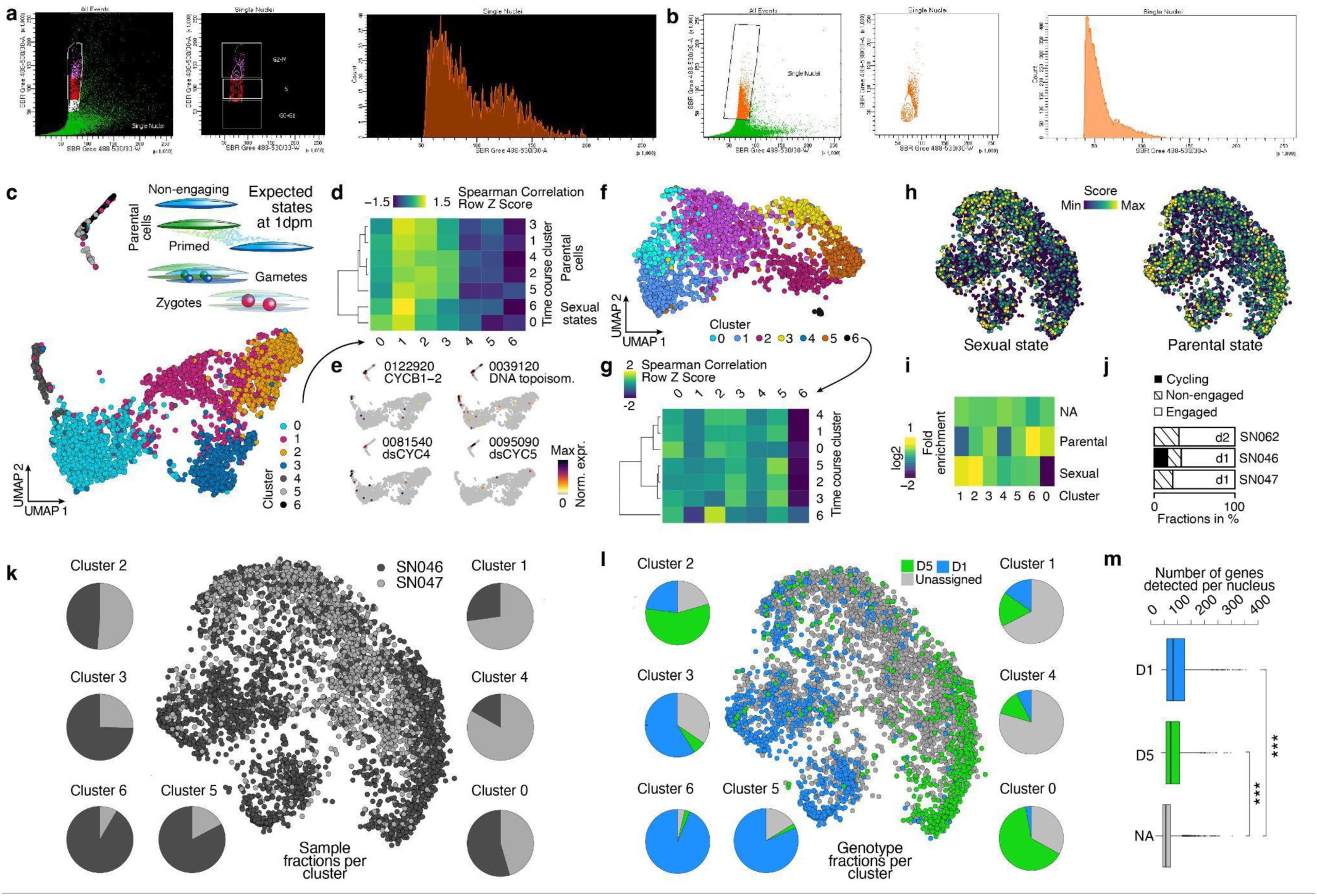
Performing snRNA-seq at the onset of sexual reproduction reveals frequent commitment of parental cells. **a-b** FACS profiles and gating strategy applied for separating single nuclei from debris after nuclei extraction for library SN046 in a and for library SN047 in b. **c** UMAP embedding of nuclei originating from library SN046. Louvain clusters are color-coded. Inset: Schematic highlights cellular states and stages expected after 1 day post mating (dpm). **d** Averaged expression per clusters of c are correlated (Spearman) with cluster averages of the time course data set (see Fig. 2b) and results are shown as Heatmap. **e** Canonical mitosis marker gene expression is projected on the UMAP of c. **f** UMAP embedding of nuclei originating from library SN047. Louvain clusters are colorcoded. **g** Averaged expression per clusters of f are correlated (Spearman) with cluster averages of the time course data set (see Fig. 2b) and results are shown as Heatmap. **h** Sexual and parental state scores are projected on the cycling cell free UMAP embedding. **i** Enrichment values of assigned states per cluster are represented as heatmap. **j** Histogram shows the relative fractions of committed, non-committed and cycling cells across 3 single-cell libraries with more than 1000 nuclei detected. **k** UMAP embedding is colored by library identity. Pie charts represent the fractions per cluster (see Fig. 3). **l** UMAP embedding is colored by genotype assignment. Pie charts represent the fractions per cluster (see Fig. 3). **m** Boxplot representation of detected number of genes across the assigned genotype identities.

**Extended Data Fig. 4.**
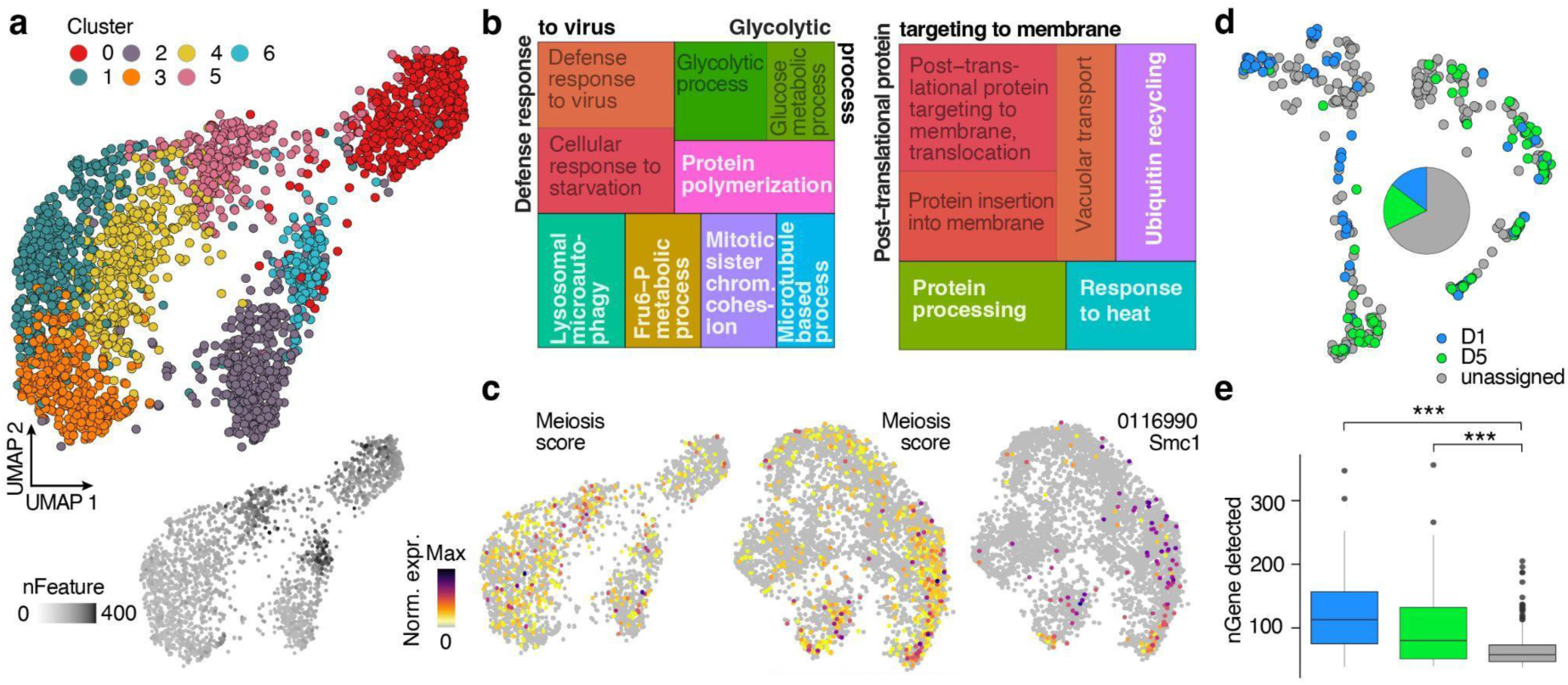
Committed cells show pre-meiotic features. **a** Committed MT-(cluster 5) and MT+ (cluster 0) and early sexual state cells (cluster 2) (see Fig. 3b) were re-clustered (Louvain) and re-embedded. The result is visualized as UMAP. Bottom: Number of detected genes is projected on the UMAP. **b** A treemap plot visualizes GO terms identified by a GO enrichment analysis on the cluster markers in a. Terms in white or outside of the boxes are the overarching parental terms for similar colored GO terms. **c** A meiosis score was calculated across all nuclei of the 1 day samples. Feature plots show score values on the gamete to zygote embedding (left), the cell cycle filtered embedding (center) and the expression level of Smc1 (right). **d** Genotype assignment for cycling cells (see Fig. 3c). Pie chart summarizes different fractions. **e** Boxplots of detected gene numbers for cells across the genotype groups. Pairwise Wilcoxon test ***: p value < 0.0001.

**Extended Data Fig. 5.**
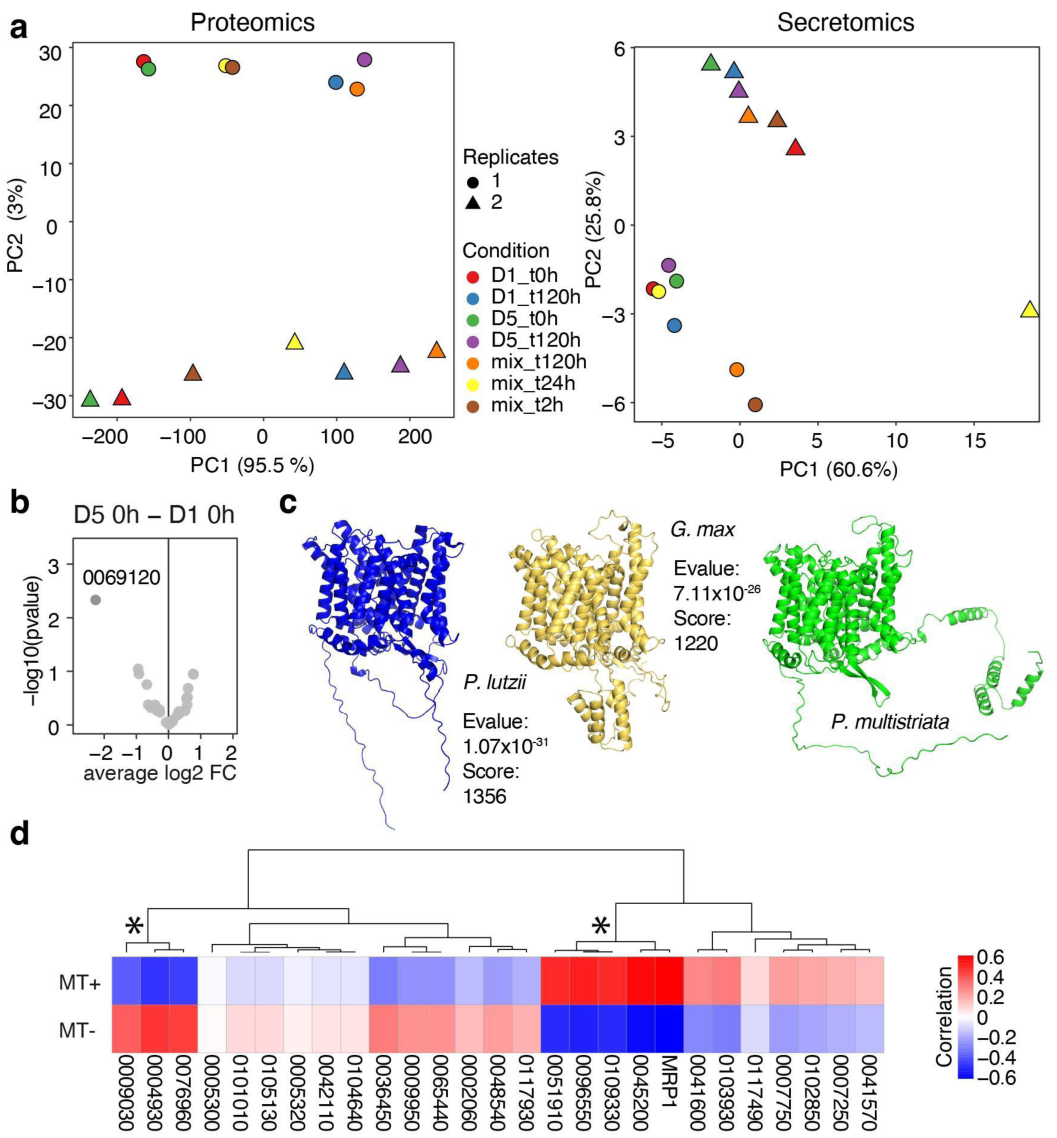
Protein structures of sex specific proteins show a high degree of conservation. **a** PCA plots describe variation between replicates of the protein abundance measurements of the whole cellular proteome (left) and the secretome (right). **b** Vulcano plot shows the differences between the 0h parental D5 and D1 secretome samples, respectively. Negative fold change values represent D5 specific proteins and positive values D1 specific proteins. Only significant hits are labelled. **c** Alphafold predictions for 3 aligned bicarbonate transporters are shown. Structural alignment scores obtained from foldseek after querying the *P. multistriata* bicarbonate transporter structure is provided. **d** Heatmap showing correlation values to pseudoMT specific markers, respectively. Genes are hierarchical clustered and asterisks mark branches with average correlation values above 0.5. Genes within those branches were screened for a matching protein profile between the MTs.

**Extended Data Fig. 6.**
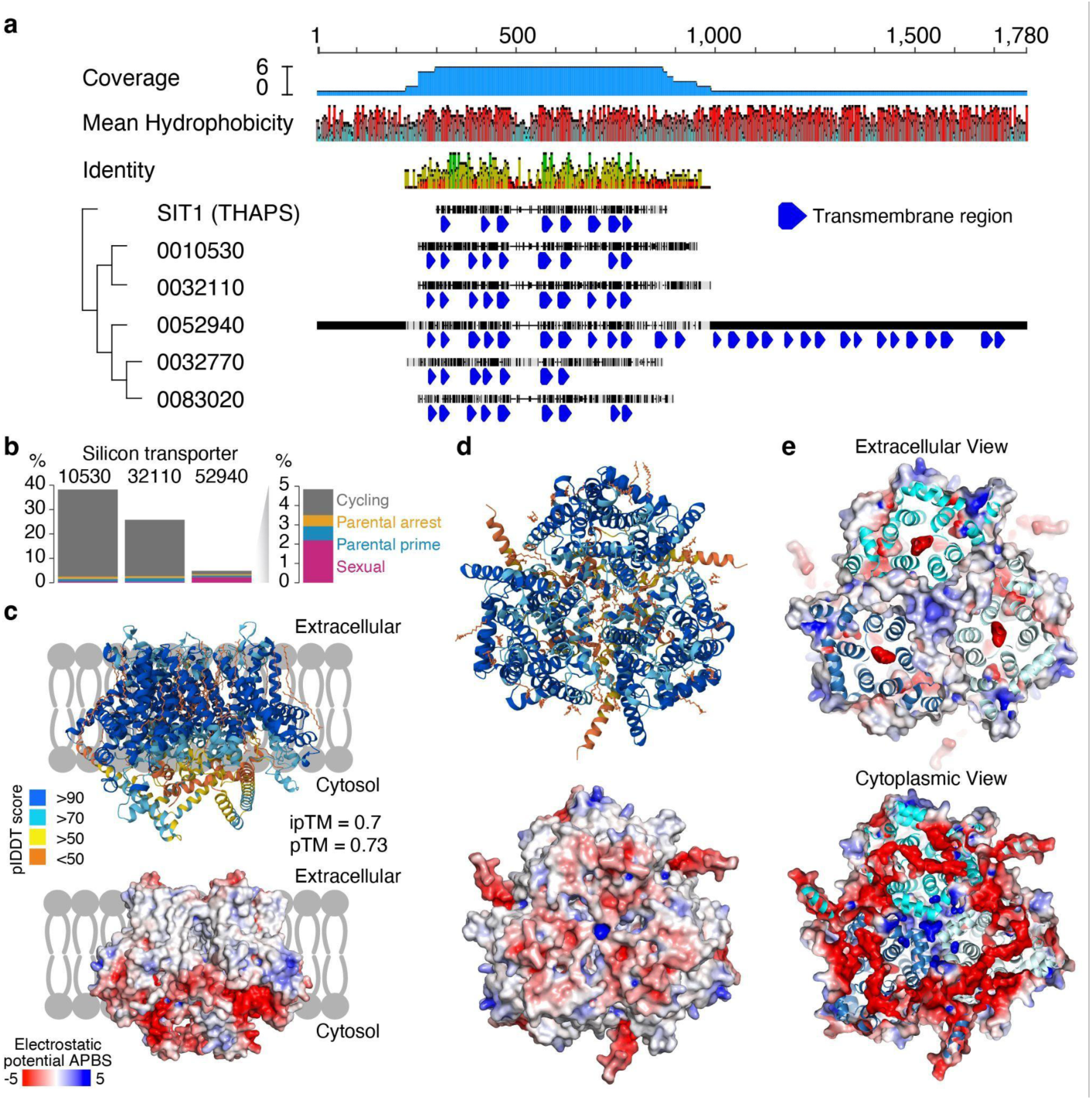
Sex and cell cycle specific usage of SITs. **a** Protein alignment of *P. multistriata* SITs identified and SIT1 of T. pseudonana. Alignment and tree are exported from GeneiousPrime. **b** Histogram with binarized expression of SITs across the cell state groupings. Note the general low detection of 52940. **c** Sideview on the Alphafold predicted homotrimer of the protein 10530. Top: Structure colored by prediction confidence. Bottom: APBS electrostatic surface model predicted by PyMOL. A gray lipid bilayer cartoon is added to the background for easier visualization. **d** Extracellular view on the predicted homotrimer. Top: Structure colored by prediction confidence. Bottom: APBS electrostatic surface model predicted by PyMOL. **e** Mixed visualization of predicted structure and APBS electrostatic surface model. Top: Extracellular view with depth fading highlights negative electrostatic charge in the center of the translocation pathways and neutral external surface areas. Bottom: Cytoplasmic view highlights a ring of negative electrostatic charge and a positively charged core between the monomers.

## Methods

### Strains, culture condition and experimental crosses

*P. multistriata* cells were grown in a f/2 Guillard medium^50^, in artificial sea water, in 12 hours light / 12 hours dark cycles at 18℃, at an irradiance of 60 µmol photons m^-2^s^-1^. Strains used in the experiment were wild strains 1265D1, 1265D5, 1264-3, 1264-2 isolated between 2020 and 2022 from the Long-Term Ecological Research station MareChiara in the Gulf of Naples (Italy) lab strains VD10 and MF4 originated from lab crosses. For nuclei capture we inoculated parental strain as monocultures at a cell density of 80,000 cells /ml.

For the time-course experiment, a unique 75 cm^2^ cell culture flask was inoculated and cells were withdrawn for each of the three time points (24, 48 and 72 hours) for nuclei extraction.

### Isolation of cellular stages by image-enabled cell sorting

Diatom parental cultures and crosses were collected, washed in ASW (artificial sea water), and incubated with Sybr Green (Thermo Fisher Scientific, Waltham, USA) diluted (1:1000) in ASW for 5 minutes. Samples were acquired and analyzed on an early-access prototype of the BD FACSDiscover™ S8 Cell Sorter^17^. The prototype was equipped with three lasers (405 nm, 488 nm, and 633 nm) and featured BD CellView™ Image Technology for image capture and processing.

Sybr Green fluorescence (DNA staining), chlorophyll autofluorescence, and selected image-derived parameters, including eccentricity, were used to establish gating strategies that enabled the identification and isolation of sexual stages such as gametes/zygotes, auxospores, and parental cells. Furthermore, the DNA-staining dye allowed the analysis of the cell cycle phases.

Cells were (50 parental cells/20 sexual state cells per well, respectively) were directly sorted into 10 µL lysis buffer (6M GuHCl, 50 mM Tris, 1% TritonX, 10 mM DTT)^51^ and samples were briefly spun down and afterwards frozen at -80℃ overnight. We collected and extracted RNA from a range of 20-50 cells for each condition, parental cells (5 replicates) and sexual subtypes (7 replicates), performed SmartSeq2^52^ and sequenced the mini-bulk transcriptomes. After thawing on ice, the suspension was homogenized by pipetting and SPRIselect beads were added at a ratio (2.2x vol:vol) to each well. A single 80% EtOH wash was performed and samples were eluted off the beads in 10µL H_2_O. 2.4 µL of the elution were used as input for SmartSeq2^52^ following a slightly modified protocol (SuperScriptIV instead of II) optimized by EMBL’s GeneCore Facility. Preamplification was done with 18 PCR cycles. Libraries were sequenced on a MiSeq with the MICRO kit reading 150bp-8-8-150bp.

### Morphology based genes expression analysis

Raw reads were mapped to the *P. multistriata* genome using STARsolo (v2.7.9)^53^ with the SmartSeq adjusted gene counting strategy. We also sorted, sequenced and mapped original parental cells but did not include these samples in the analysis. An initial screening revealed a few samples with only few detected genes per sample and based on this finding we removed samples with less than 300 detected genes per sample. This left us with 12 samples in total for data analysis. These remaining samples were downsampled to 5000 counts since 2 samples had a disproportionally high-count value compared to the remaining samples. Next, we log normalized the counts, scaled the data with regressing out the number of counts and ran a PCA in Seurat. Marker genes that were used as input for scoring the single-cell data were obtained by using Seurat’s FindMarkers() function by comparing the 2 states against each other (Pos FC = parental specific, Neg FC = sex specific).

### Single Nuclei Extraction, FACS Enrichment, Capture and Library Preparation

Nuclei were extracted based on the protocol described by^54^. Briefly, cells were counted and collected by centrifugation, then permeabilized using a mild acid solution. The samples were subsequently subjected to sonication to disrupt the cells and release nuclei. After extraction, nuclei were incubated with Sybr Green (1:1000) in the NIB buffer for 5 minutes. Stained samples were analyzed on a BD FACS Aria Fusion equipped with five lasers at 355 nm, 405 nm, 488 nm, 561 nm, and 633 nm. Sybr Green was excited with the 488 nm blue laser, and emission was collected using a 530/30 band-pass filter. Sybr Green fluorescence was used to label the nuclei gating out debris and aggregates. Depending on the sample, between 200.000 and 500.000 nuclei were bulk-sorted in purity mode per condition.

Following FACS enrichment, nuclei quality and concentration were assessed by staining with SYBR Safe DNA stain and imaging on a Zeiss Axio Imager using a C-Chip hemocytometer (Neubauer improved). Single nuclei capture and library preparation were performed using the Chromium Single Cell 3’ Gene Expression Kit (v3.1 chemistry) and the Chromium Controller (10x Genomics), following the manufacturer’s protocol with slight modifications. Approximately 20,000 nuclei were loaded per capture channel. cDNA amplification was performed with 12 cycles, using a 3-minute extension time. Final libraries were quantified using a Qubit fluorometer, and fragment sizes were assessed using an Agilent Bioanalyzer. Libraries were sequenced with 250 million PE reads on an Illumina NextSeq 2000 using the following read configuration: 28 cycles for Read 1, 10 cycles each for the i5 and i7 indices, and 90 cycles for Read 2.

### Experimental design for 10x runs

The RNA content of *P. multistriata* nuclei is very low. Initial trials with loading nuclei suspensions on the 10x Genomics platform using the Chromium Single Cell 3′ v3 kit resulted in no satisfying results after cDNA pre-amplication and in some cases an excess in amplification artefacts. To avoid the generation of artefacts due to low RNA input and as a positive side effect increase the number of nuclei that can be loaded on a single 10X lane we designed species mixing experiments of up to 3 species mixed. We aimed to load 20,000 diatom nuclei per 10X lane and adjusted the number of nuclei of the other species such that a total of 30,000 to 40,000 nuclei depending on the number of species mixed were loaded. This strategy increased the yield of cDNA preamplification by eliminating artefacts. Results were comparable across all species mixing experiments performed.

Experiments of the time course (SN061/62/63) were carried out in parallel using the same cDNA generation mastermix to reduce batch effects. To achieve this, all nuclei had been frozen after they had been isolated and sorted. The same strategy was used for the 1day samples. However, only one sample (SN046) was frozen prior to loading while the other sample (SN047) was directly loaded after sorting which resulted in a substantial batch effect even though the same cDNA master mix was used. Interestingly, the batch effect is not affecting the general QC statistics (Extended Data Fig. 1).

### Data processing

Raw 10x reads were mapped to concatenated pan genomes of the respective species using Cellranger v6 (10X Genomics). Genomes were retrieved from EnsemblMetazoa (https://metazoa.ensembl.org) or EnsemblProtists (https://protists.ensembl.org) and species mixes included in addition to *P. multistriata* either *B. lanceolatum* or *P. dumerilii* or both (Extended Data Table 1).

Raw RNA count results were loaded into Seurat with only very mild filtering criteria (min.cells = 2, min.features = 100, nCounts < 30,000). The count information of genes of the different species was used to calculate a species score by counting the number of genes expressed for a certain species normalized to the total number of genes in the respective genome and subsequent transformation to a 0 to 1 scale. The score was used to determine species doublets (all species scores > 0.05) and empty droplets with very low scores for any of the species (all species scores < 0.05) which were removed from the data, respectively. Afterwards, the default Seurat pipeline (Normalization > Scaling > PCA > Louvain clustering > UMAP embedding) was used to determine and visualize clusters that were classified by species origin based on the species score. The clustering revealed in all data sets a clear separation by species which was used to subset the data into species specific count matrices. Non *P. multistriata* genes were removed from the diatom data sets and all analyses presented in this study originate from these cleaned diatom specific matrices.

### Data analysis

All basic single-cell data processing steps were performed in R using the R package Seurat (v4)^55^. Briefly, count matrices of 10x and SmartSeq2 experiments were log normalized and scaling was performed across all genes with regressing out nCounts. An initial round of processing the data sets revealed clusters that were characterized by low counts and almost to no marker genes per cluster. After removing such outlier clusters, the process was repeated. Similarly, samples of the SmartSeq2 experiments were filtered For the time course data set 15 PCs were used to cluster and embed the nuclei by Uniform Manifold Approximation and Projection (UMAP) to reduce the dimensionality for data exploration. The 1 day data sets showed a remarkable batch effect between libraries and an integration by Harmony^56^ using top 50 PCs was needed. Afterwards, 20 Harmony components were used to cluster (Louvain) and embed (UMAP) the nuclei.

Scores like the Meiosis score were calculated using the AddModuleScore() function in Seurat using default parameters and custom gene sets as input. For scoring the single-cell data based on the phenotype sorted data, top 50 state specific markers were used. Note that the markers were determined based on a non-downsampled data set to not risk losing important genes that might have been lost during downsampling.

Pseudotime estimates were calculated using the R package ‘destiny’^57^ or the R package ‘Monocle3’^58^

Heatmap visualizations were generated using the R package ‘pheatmap’

Protein alignments and tree estimates were performed using Geneious Prime and its internal Geneious Prime Alignment software with a global alignment type with free end gaps (gap open penalty= 3/ Gap extension penalty = 1).

GO enrichment analyses were performed using GoSeq^59^ and were visualized using the R package rrvgo

For all other visualization and plot types basic R functions and the R package ggplot2 were used.^60^

### Protein predictions

The Alphafold server (https://alphafoldserver.com/) was used to model protein structures based on amino acid sequences retrieved from Uniprot (www.uniprot.org) in case there were no predicted structures provided. In case of missing amino acid information in the protein sequences (indicated by X) the corresponding positions were replaced by Alanines (A) in order to allow a prediction using Alphafold. When predicting the protein assembly of the SIT monomer (10530) we added 50 oleic acid ligands (OLA) to the prediction to mimic the presence of a lipid bi-layered membrane. For all protein structure visualization, we used the software PyMOL (v3.0.5).

### Proteomics sample preparation

Diatom samples from a time course were lysed according to the lysis protocol described (PMID: 39930009). Generated tryptic peptides were labeled using TMT10plex™ reagent as previously described (PMID: 24579773). Labeled samples were combined for multiplexing, desalted using an Oasis® HLB µElution Plate (Waters) according to the manufacturer’s instructions, and dried by vacuum centrifugation.

Offline high-pH reversed-phase fractionation (PMID: 22462785) was carried out using an Agilent 1200 Infinity high-performance liquid chromatography (HPLC) system, equipped with a Gemini C18 analytical column (3 μm particle size, 110 Å pore size, dimensions 100 x 1.0 mm, Phenomenex) and a Gemini C18 SecurityGuard pre-column cartridge (4 x 2.0 mm, Phenomenex). The mobile phases consisted of 20 mM ammonium formate adjusted to pH 10.0 (Buffer A) and 100% acetonitrile (Buffer B). The peptides were separated at a flow rate of 0.1 mL/min using the following linear gradient: 100% Buffer A for 2 minutes, ramping to 35% Buffer B over 59 minutes, increasing rapidly to 85% Buffer B within 1 minute, and holding at 85% Buffer B for an additional 15 minutes. Subsequently, the column was returned to 100% Buffer A and re-equilibrated for 13 minutes. During the LC separation, 48 fractions were collected. These were pooled into twelve fractions by combining every twelfth fraction. The pooled fractions were then dried using vacuum centrifugation.

An UltiMate 3000 RSLCnano LC system (Thermo Fisher Scientific) equipped with a trapping cartridge (µ-Precolumn C18 PepMap™ 100, 300 µm i.d. × 5 mm, 5 µm particle size, 100 Å pore size; Thermo Fisher Scientific) and an analytical column (nanoEase™ M/Z HSS T3, 75 µm i.d. × 250 mm, 1.8 µm particle size, 100 Å pore size; Waters). Samples were trapped at a constant flow rate of 30 µL/min using 0.05% trifluoroacetic acid (TFA) in water for 6 minutes. After switching in-line with the analytical column, which was pre-equilibrated with solvent A (3% dimethyl sulfoxide [DMSO], 0.1% formic acid in water), the peptides were eluted at a constant flow rate of 0.3 µL/min using a gradient of increasing solvent B concentration (3% DMSO, 0.1% formic acid in acetonitrile). The gradient was as follows: 2% to 8% in 4 minutes (min), 8% to 28% in 104 min, 28% to 40% in 4 min, 40%-80% in 4 min and re-equilibrated to 2% B for 4 min. Peptides were introduced into a Q Exactive™ Plus mass spectrometer (Thermo Fisher Scientific) via a Pico-Tip emitter (360 µm OD × 20 µm ID; 10 µm tip, CoAnn Technologies) using an applied spray voltage of 2.2 kV. The capillary temperature was maintained at 275 °C. Full MS scans were acquired in profile mode over an m/z range of 375–1,200, with a resolution of 70,000 at m/z 200 in the Orbitrap. The maximum injection time was set to 250 ms, and the AGC target limit was set to 3E6. The instrument was operated in data-dependent acquisition (DDA) mode, with MS/MS scans acquired in the Orbitrap at a resolution of 35,000. The maximum injection time was set to 120 ms, with an AGC target of 2E5. Fragmentation was performed using higher-energy collisional dissociation (HCD) with a normalized collision energy of 32%, and MS2 spectra were acquired in profile mode. The quadrupole isolation window was set to 0.7 m/z, and dynamic exclusion was enabled with a duration of 30 seconds. The peptide match algorithm was set to ‘preferred’ and charge exclusion ‘unassigned’. Only precursor ions with charge states 2–4 were selected for fragmentation.

Description of PRIDE upload. Mention which information to put and to share reviewer login details.

### Secretory proteomics sample preparation

Proteins were precipitated using ammonium sulfate to saturation. Briefly, 0.71 g of ammonium sulfate was added to 1 mL aliquots of protein sample and incubated overnight at 4 °C with gentle rotation. The following day, samples were rotated at room temperature for 30 min to partially dissolve residual salt, then centrifuged at 16,000 × g for 45 min at 0 °C. The supernatant was removed, and the resulting protein pellet was resuspended in 50 µL of 1% SDS. Samples were incubated for 1 h at 50 °C and 1,200 rpm to ensure complete solubilization of the protein pellet.

Protein digestion was performed using the Single-Pot Solid-Phase-enhanced Sample Preparation (SP2) method. Each solubilized sample (approximately 100 µL total volume) was combined with 150 µL ethanol, 2 µL magnetic beads, and 50 µL of 15% formic acid. Samples were incubated at room temperature for 15 min with shaking. Following incubation, two phases were typically observed—an upper phase containing magnetic beads and a lower phase containing salt precipitate. The beads were collected on a magnetic stand and washed twice with 70% ethanol and once with acetonitrile (ACN). Proteins bound to the beads were digested overnight with trypsin at room temperature. The next day, samples were dried completely prior to labeling. Following labeling, samples were desalted using OASIS cartridges according to the manufacturer’s protocol.

Desalted samples were dried, reconstituted, and 10% of each sample was injected into a Q Exactive mass spectrometer (Thermo Fisher Scientific) for LC–MS/MS analysis.

Dried peptide samples were resuspended in 10 µL of water and labeled with 4 µL of TMT10plex reagent according to the manufacturer’s instructions.

### Proteomics data processing

Raw files were converted to mzML format using MSConvert from ProteoWizard, using peak picking, 64-bit encoding and zlib compression, and filtering for the 1000 most intense peaks. Files were then searched using MSFragger in FragPipe (unknown) against FASTA database uniprot-author ferrante+i+m -filtered-reviewed no containing common contaminants and reversed sequences. The following modifications were included into the search parameters: Carbamidomethylation (C, 57.0215), TMT (K, 229.1629) as fixed modifications; Oxidation (M, 15.9949), Acetylation (protein N-terminus, 42.0106), TMT (peptide N-terminus, 229.1629) as variable modifications. For the full scan (MS1) a mass error tolerance of 20 PPM and for MS/MS (MS2) spectra of 20 PPM was set. For protein digestion, ’trypsin’ was used as protease with an allowance of maximum 2 missed cleavages requiring a minimum peptide length of 7 amino acids. The false discovery rate on peptide and protein level was set to 0.01. The standard settings of the FragPipe workflow ’TMT10’ were used. The following modifications were made: msfragger.add_topN_complementary: 0, msfragger.misc.fragger.enzyme-dropdown-1: trypsin, msfragger.misc.fragger.precursor-charge-hi: 6, msfragger.search_enzyme_name_1: trypsin, msfragger.search_enzyme_nocut_1: P, msfragger.use_topN_peaks: 300, msfragger.write_calibrated_mgf: false, peptide-prophet.run-peptide-prophet: true, tmtintegrator.allow_unlabeled: true, tmtintegrator.channel_num: 6, tmtintegrator.dont-run-fq-lq: false, tmtintegrator.unique_gene: 1, tmtintegrator.unique_pep: true.

### Proteomics data analysis

For the proteomics data analysis, the raw output files of FragPipe (protein.tsv files files) were processed using the R programming environment (ISBN 3-900051-07-0). Initial data processing included filtering out contaminants and reverse proteins. Only proteins quantified with at least 2 razor peptides (with Razor.Peptides >= 2) were considered for further analysis. 3593 proteins passed the quality control filters. In order to correct for technical variability, batch effects were removed using the ’removeBatchEffect’ function of the limma package (PMID: 25605792) on the log2 transformed raw TMT reporter ion intensities (’channel’ columns). Subsequently, normalization was performed using the ’normalizeVSN’ function of the limma package (VSN - variance stabilization normalization - PMID: 12169536). Missing values were imputed with the ’knn’ method using the ’impute’ function from the Msnbase package (PMID: 22113085). This method estimates missing data points based on similarity to neighboring data points, ensuring that incomplete data did not distort the analysis. Differential expression analysis was performed using the moderated t-test provided by the limma package (PMID: 25605792). The model accounted for replicate information by including it as a factor in the design matrix passed to the ’lmFit’ function. Imputed values were assigned a weight of 0.01 in the model, while quantified values were given a weight of 1, ensuring that the statistical analysis reflected the uncertainty in imputed data. Proteins were annotated as hits if they had a false discovery rate (FDR) below 0.05 and an absolute fold change greater than 2. Proteins were considered candidates if they had an FDR below 0.2 and an absolute fold change greater than 1.5. Clustering with all hit proteins based on the median protein abundances normalized by median of control condition was conducted to identify groups of proteins with similar patterns across conditions. The ’kmeans’ method was employed, using Euclidean distance as the distance metric and ’ward.D2’ linkage for hierarchical clustering. The optimal number of clusters (16) was determined using the Elbow method, which identifies the point where the within-group sum of squares stabilizes.

## Extended Data Files

**Extended Data Table 1.** SnRNA-seq experimental overview and summary statistics.

| Library ID | Time after crossing | Freezing* | Average nCells | Average nGenes | Additional species |
| --- | --- | --- | --- | --- | --- |
| SN046 | 1 day | no | 3242 | 77 | *P. dumerilii* |
| SN047 | 1 day | yes | 2131 | 70 | *P. dumerilii* |
| SN061 | 1 day | yes | 179 | 118 | *B. lanceolatum* |
| SN062 | 2 day | yes | 1041 | 136 | *P. dumerilii/B. lanceolatum* |
| SN063 | 3 day | yes | 288 | 132 | *P. dumerilii/B. lanceolatum* |

\* Additional round of freezing at -80 after nuclei extraction

**Extended Data Table 2.** Diatom proteomics reveals protein differences across a time course of sexual reproduction. Hit: significant, Candidate: reduced significance, No hit: non-significant.

| Comparison | Hit | Candidate | No hit |
| --- | --- | --- | --- |
| (D5_t120h-D1_t120h)-(D5_t0h-D1_t0h) | 0 | 0 | 3593 |
| (D5_t120h-mix_t120h)-(D1_t120h-mix_t120h) | 9 | 40 | 3544 |
| D1_t120h-D1_t0h | 945 | 939 | 1709 |
| D1_t120h-mix_t120h | 0 | 4 | 3589 |
| D5_t0h-D1_t0h | 7 | 13 | 3573 |
| D5_t120h-D1_t120h | 9 | 40 | 3544 |
| D5_t120h-D5_t0h | 223 | 1036 | 2334 |
| D5_t120h-mix_t120h | 0 | 6 | 3587 |
| mix_t120h-D1_t0h | 858 | 968 | 1767 |
| mix_t120h-D5_t0h | 548 | 929 | 2116 |
| mix_t120h-mix_t24h | 1 | 131 | 3461 |
| mix_t120h-mix_t2h | 0 | 60 | 3533 |
| mix_t24h-D1_t0h | 6 | 289 | 3298 |
| mix_t24h-D5_t0h | 2 | 0 | 3591 |
| mix_t24h-mix_t2h | 2 | 0 | 3591 |
| mix_t2h-D1_t0h | 0 | 0 | 3593 |
| mix_t2h-D5_t0h | 0 | 1 | 3592 |

**Extended Data Table 3.** Diatom secretomics reveals secreted protein differences across a time course of sexual reproduction. Hit: significant, Candidate: reduced significance, No hit: non-significant.

| Comparison | Hit | Candidate | No hit |
| --- | --- | --- | --- |
| (D5_t120h-D1_t120h)-(D5_t0h-D1_t0h) | 0 | 0 | 29 |
| (D5_t120h-mix_t120h)-(D1_t120h-mix_t120h) | 1 | 7 | 21 |
| D1_t120h-D1_t0h | 0 | 0 | 29 |
| D1_t120h-mix_t120h | 0 | 0 | 29 |
| D5_t0h-D1_t0h | 0 | 1 | 28 |
| D5_t120h-D1_t120h | 0 | 0 | 29 |
| D5_t120h-D5_t0h | 0 | 0 | 29 |
| D5_t120h-mix_t120h | 0 | 0 | 29 |
| mix_t120h-D1_t0h | 1 | 3 | 25 |
| mix_t120h-D5_t0h | 0 | 1 | 28 |
| mix_t120h-mix_t24h | 0 | 1 | 28 |
| mix_t120h-mix_t2h | 0 | 0 | 29 |
| mix_t24h-D1_t0h | 0 | 4 | 25 |
| mix_t24h-D5_t0h | 0 | 0 | 29 |
| mix_t24h-mix_t2h | 0 | 3 | 26 |
| mix_t2h-D1_t0h | 0 | 2 | 27 |
| mix_t2h-D5_t0h | 0 | 0 | 29 |

## Bibliography

1. Burki, F., Sandin, M. M. & Jamy, M. Diversity and ecology of protists revealed by metabarcoding. Current Biology 31, R1267–R1280 (2021).

2. Frada, M., Probert, I., Allen, M. J., Wilson, W. H. & De Vargas, C. The “Cheshire Cat” escape strategy of the coccolithophore *Emiliania huxleyi* in response to viral infection. Proc. Natl. Acad. Sci. U.S.A. 105, 15944–15949 (2008).

3. Von Dassow, P. et al. Life-cycle modification in open oceans accounts for genome variability in a cosmopolitan phytoplankton. The ISME Journal 9, 1365–1377 (2015).

4. Rynearson, T. A. et al. Major contribution of diatom resting spores to vertical flux in the sub-polar North Atlantic. Deep Sea Research Part I: Oceanographic Research Papers 82, 60–71 (2013).

5. Ellegaard, M. & Ribeiro, S. The long-term persistence of phytoplankton resting stages in aquatic ‘seed banks’. Biological Reviews 93, 166–183 (2018).

6. Brosnahan, M. L. et al. Rapid growth and concerted sexual transitions by a bloom of the harmful dinoflagellate *Alexandrium fundyense* (Dinophyceae). Limnology & Oceanography 60, 2059–2078 (2015).

7. Annunziata, R. et al. Trade-off between sex and growth in diatoms: Molecular mechanisms and demographic implications. Sci. Adv. 8, eabj9466 (2022).

8. Selander, E., Jakobsen, H. H., Lombard, F. & Kiørboe, T. Grazer cues induce stealth behavior in marine dinoflagellates. Proc. Natl. Acad. Sci. U.S.A. 108, 4030–4034 (2011).

9. Hamm, C. E. Architecture, ecology and biogeochemistry of Phaeocystis colonies. Journal of Sea Research 43, 307–315 (2000).

10. Benoiston, A.-S. et al. The evolution of diatoms and their biogeochemical functions. Phil. Trans. R. Soc. B 372, 20160397 (2017).

11. Bilcke, G. et al. Life Cycle Regulation. in The Molecular Life of Diatoms (eds Falciatore, A. & Mock, T.) 205–228 (Springer International Publishing, Cham, 2022). doi:10.1007/978-3-030-92499-7_8.

12. Montresor, M., Vitale, L., D’Alelio, D. & Ferrante, M. I. Sex in marine planktonic diatoms: insights and challenges. pip 3, 61–75 (2016).

13. Fromm, A. et al. Single-cell RNA-seq of the rare virosphere reveals the native hosts of giant viruses in the marine environment. Nat Microbiol 9, 1619–1629 (2024).

14. Ferrante, M. I., Broccoli, A. & Montresor, M. The pennate diatom *Pseudo-nitzschia multistriata* as a model for diatom life cycles, from the laboratory to the sea. Journal of Phycology 59, 637–643 (2023).

15. Scalco, E., Stec, K., Iudicone, D., Ferrante, M. I. & Montresor, M. The dynamics of sexual phase in the marine diatom *P seudo-nitzschia multistriata* (B acillariophyceae). Journal of Phycology 50, 817–828 (2014).

16. Marotta, P. et al. Mate Perception and Gene Networks Regulating the Early Phase of Sex in Pseudo-nitzschia multistriata. JMSE 10, 1941 (2022).

17. Schraivogel, D. et al. High-speed fluorescence image–enabled cell sorting. Science 375, 315–320 (2022).

18. Scalco, E., Amato, A., Ferrante, M. I. & Montresor, M. The sexual phase of the diatom Pseudo-nitzschia multistriata: cytological and time-lapse cinematography characterization. Protoplasma 253, 1421–1431 (2016).

19. Rizos, I., Frada, M. J., Bittner, L. & Not, F. Life cycle strategies in free-living unicellular eukaryotes: Diversity, evolution, and current molecular tools to unravel the private life of microorganisms. J Eukaryotic Microbiology 71, e13052 (2024).

20. Ning, J. et al. Comparative genomics in *Chlamydomonas* and *Plasmodium* identifies an ancient nuclear envelope protein family essential for sexual reproduction in protists, fungi, plants, and vertebrates. Genes Dev. 27, 1198–1215 (2013).

21. Russo, M. T. et al. MRP3 is a sex determining gene in the diatom Pseudo-nitzschia multistriata. Nat Commun 9, 5050 (2018).

22. Wong, J. L., Créton, R. & Wessel, G. M. The Oxidative Burst at Fertilization Is Dependent upon Activation of the Dual Oxidase Udx1. Developmental Cell 7, 801–814 (2004).

23. Patil, S. et al. Identification of the meiotic toolkit in diatoms and exploration of meiosis-specific SPO11 and RAD51 homologs in the sexual species Pseudo-nitzschia multistriata and Seminavis robusta. BMC Genomics 16, 930 (2015).

24. Haghverdi, L., Büttner, M., Wolf, F. A., Buettner, F. & Theis, F. J. Diffusion pseudotime robustly reconstructs lineage branching. Nat Methods 13, 845–848 (2016).

25. Bilcke, G. et al. Mating type specific transcriptomic response to sex inducing pheromone in the pennate diatom *Seminavis robusta*. The ISME Journal 15, 562–576 (2021).

26. Veluchamy, A. et al. Insights into the role of DNA methylation in diatoms by genome-wide profiling in Phaeodactylum tricornutum. Nat Commun 4, 2091 (2013).

27. Basu, S. et al. Finding a partner in the ocean: molecular and evolutionary bases of the response to sexual cues in a planktonic diatom. New Phytologist 215, 140–156 (2017).

28. Huysman, M. J. et al. Genome-wide analysis of the diatom cell cycle unveils a novel type of cyclins involved in environmental signaling. Genome Biol 11, R17 (2010).

29. Mager, S. et al. Genomic diversity in time and space in the toxic diatom Pseudo-nitzschia multistriata. Harmful Algae 142, 102791 (2025).

30. Li, H. A statistical framework for SNP calling, mutation discovery, association mapping and population genetical parameter estimation from sequencing data. Bioinformatics 27, 2987–2993 (2011).

31. Huang, Y., McCarthy, D. J. & Stegle, O. Vireo: Bayesian demultiplexing of pooled single-cell RNA-seq data without genotype reference. Genome Biol 20, 273 (2019).

32. Bilcke, G. et al. Conserved genetic markers reveal widespread diatom sexual reproduction in the global ocean. Nat Commun 16, 10029 (2025).

33. Lemonnier, T., Dupré, A. & Jessus, C. The G2-to-M transition from a phosphatase perspective: a new vision of the meiotic division. Cell Div 15, 9 (2020).

34. Kar, F. M. & Hochwagen, A. Phospho-Regulation of Meiotic Prophase. Front. Cell Dev. Biol. 9, 667073 (2021).

35. Knecht, S. et al. An Introduction to Analytical Challenges, Approaches, and Applications in Mass Spectrometry–Based Secretomics. Molecular & Cellular Proteomics 22, 100636 (2023).

36. Bücherl, C. A. et al. Plant immune and growth receptors share common signalling components but localise to distinct plasma membrane nanodomains. eLife 6, e25114 (2017).

37. Ruggiero, A. et al. DNA methylation is linked to the monoallelic expression of *MRP3*, a diatom mating type determining gene. Preprint at 10.1101/2023.10.11.561864 (2023).

38. Shrestha, R. P. & Hildebrand, M. Evidence for a Regulatory Role of Diatom Silicon Transporters in Cellular Silicon Responses. Eukaryot Cell 14, 29–40 (2015).

39. Knight, M. J., Senior, L., Nancolas, B., Ratcliffe, S. & Curnow, P. Direct evidence of the molecular basis for biological silicon transport. Nat Commun 7, 11926 (2016).

40. Villar, E. et al. DIATOMICBASE : a versatile gene-centered platform for mining functional omics data in diatom research. The Plant Journal 121, e70061 (2025).

41. Durkin, C. A., Koester, J. A., Bender, S. J. & Armbrust, E. V. The evolution of silicon transporters in diatoms. Journal of Phycology 52, 716–731 (2016).

42. Alverson, A. J. et al. Phylogenomics reveals the slow-burning fuse of diatom evolution. Proc. Natl. Acad. Sci. U.S.A. 122, e2500153122 (2025).

43. Honigberg, S. M. & Purnapatre, K. Signal pathway integration in the switch from the mitotic cell cycle to meiosis in yeast. Journal of Cell Science 116, 2137–2147 (2003).

44. Van Werven, F. J. & Amon, A. Regulation of entry into gametogenesis. Phil. Trans. R. Soc. B 366, 3521–3531 (2011).

45. Font-Muñoz, J. S. et al. Collective sinking promotes selective cell pairing in planktonic pennate diatoms. Proc. Natl. Acad. Sci. U.S.A. 116, 15997–16002 (2019).

46. MacKenzie, A., Vicory, V. & Lacefield, S. Meiotic cells escape prolonged spindle checkpoint activity through kinetochore silencing and slippage. PLoS Genet 19, e1010707 (2023).

47. Flori, S. et al. Diatom ultrastructural diversity across controlled and natural environments. Preprint at 10.1101/2025.06.16.659906 (2025).

48. Russo, M. T., Aiese Cigliano, R., Sanseverino, W. & Ferrante, M. I. Assessment of genomic changes in a CRISPR/Cas9 *Phaeodactylum tricornutum* mutant through whole genome resequencing. PeerJ 6, e5507 (2018).

49. Yin, W. & Hu, H. CRISPR/Cas9-Mediated Genome Editing via Homologous Recombination in a Centric Diatom *Chaetoceros muelleri*. ACS Synth. Biol. 12, 1287–1296 (2023).

50. Guillard, R. R. L. Culture of Phytoplankton for Feeding Marine Invertebrates. in Culture of Marine Invertebrate Animals (eds Smith, W. L. & Chanley, M. H.) 29–60 (Springer US, Boston, MA, 1975). doi:10.1007/978-1-4615-8714-9_3.

51. Wollny, D., Zhao, S. & Martin-Villalba, A. RNA-seq library preparation from single pancreatic acinar cells. Preprint at 10.1101/085696 (2016).

52. Picelli, S. et al. Full-length RNA-seq from single cells using Smart-seq2. Nat Protoc 9, 171–181 (2014).

53. Kaminow, B., Yunusov, D. & Dobin, A. STARsolo: accurate, fast and versatile mapping/quantification of single-cell and single-nucleus RNA-seq data. Preprint at 10.1101/2021.05.05.442755 (2021).

54. Annunziata, R. et al. An optimised method for intact nuclei isolation from diatoms. Sci Rep 11, 1681 (2021).

55. Hao, Y. et al. Integrated analysis of multimodal single-cell data. Cell 184, 3573–3587.e29 (2021).

56. Korsunsky, I. et al. Fast, sensitive and accurate integration of single-cell data with Harmony. Nat Methods 16, 1289–1296 (2019).

57. Angerer, P., et al. *destiny* – diffusion maps for large-scale single-cell data in R. Preprint at 10.1101/023309 (2015).

58. Trapnell, C. et al. The dynamics and regulators of cell fate decisions are revealed by pseudotemporal ordering of single cells. Nat Biotechnol 32, 381–386 (2014).

59. Young, M. D., Wakefield, M. J., Smyth, G. K. & Oshlack, A. Gene ontology analysis for RNA-seq: accounting for selection bias. Genome Biol 11, R14 (2010).

60. Wickham, H. Ggplot2: Elegant Graphics for Data Analysis. (Springer international publishing, Cham, 2016).

